# BunDLe-Net: Learning minimal neural manifolds that represent only behaviourally relevant dynamics

**DOI:** 10.1101/2023.08.08.551978

**Authors:** Akshey Kumar, Aditya Gilra, Anja Meunier, Moritz Grosse-Wentrup

**Affiliations:** Research Group Neuroinformatics, Faculty of Computer Science, University of Vienna, Austria; Machine Learning Group, Centrum Wiskunde & Informatica, Amsterdam, Netherlands; Department of Computer Science, The University of Sheffield, Sheffield, United Kingdom; UniVie Doctoral School Computer Science DoCS, University of Vienna, Austria; Vienna Cognitive Science Hub, Vienna, Austria; Data Science @ Uni Vienna, Vienna, Austria; Wirtschaftsuniversität, Vienna, Austria

## Abstract

Neural manifold learning techniques represent high-dimensional neural dynamics in low-dimensional embeddings to reveal the intrinsic structure of neural manifolds. We introduce a novel neural manifold learning technique, BunDLe-Net, that learns a low-dimensional Markovian embedding of the neural dynamics which preserves only those aspects of the neural dynamics that are relevant for a given behavioural context. In contrast to other techniques that learn embeddings which allow a good reconstruction of the original data, BunDLe-Net eliminates neural dynamics that are irrelevant for decoding behaviour, effectively de-noising the data to reveal better the intricate relationships between neural dynamics and behaviour. We show that BunDLe-Net learns highly consistent manifolds across animals that reveal the building blocks of their neural manifolds on a variety of data sets, ranging from calcium imaging data recorded in the nematode *C. elegans* to spiking data from the rat hippocampus.

## Introduction

Neural manifold learning (NML) has emerged as a powerful tool to analyse high-dimensional brain recordings (Mitchell-Heggs et al., 2023; Cunningham and Yu, 2014a; Fortunato et al., 2024; Vyas et al., 2020; Ebitz and Hayden, 2021; Pei et al., 2021; Duncker and Sahani, 2021; Pandarinath et al., 2018a). They are typically evaluated on how well they preserve information in the low-dimensional embeddings about the original neural dynamics and the experimental conditions, e.g., as measured by their ability to reconstruct the original neural activity from the manifold (Pei et al., 2021; Pandarinath et al., 2018b) and their performance in decoding visual stimuli (Schneider et al., 2023) or behaviours (Sani et al., 2024). However, it remains a topic of debate in which sense the low-dimensional representations provide new insights into the structure of neural dynamics (Humphries, 2020; Dyer and Kording, 2023; Perich, 2024; Saxe et al., 2021).

In this work, we argue that NML techniques should be designed from first principles to yield insights into the structure of neural dynamics. To accomplish this goal, we introduce a framework for learning low-dimensional Markovian neural representations that abstract away all dynamics in the neural recordings that are not required to explain a particular set of behaviours. We prove theoretically that our approach gives rise to neural manifolds that support causal statements about neural dynamics and demonstrate experimentally that the learned manifolds reveal how the structure of neural dynamics coordinates behavioural flexibility.

Various reconstruction-based techniques have been used for NML, such as conventional dimensionality reduction methods (PCA, t-SNE, UMAP) (Tombaz et al., 2020; Dimitriadis et al., 2018a), as well as methods explicitly designed for NML. Notable examples are PCA-based approaches like j-PCA (Churchland et al., 2012) and demixed-PCA (Kobak et al., 2016a) and autoencoder-based methods such as LFADS (Pandarinath et al., 2018b), pi-VAE (Zhou and Wei, 2020a), and distill-VAE (Li et al., 2024). While these methods excel at data compression, they may retain information in the neural manifold that is irrelevant in a given behavioural context, potentially occluding the neural dynamics that coordinate behavioural flexibility (Doerig et al., 2023). For instance, when studying an animal’s ability to navigate a maze using visual cues, neural activity patterns carrying auditory or olfactory information may be irrelevant to the behavioural context and could be abstracted away to better reveal relationships between neural representations of the visual cues and motor behaviour. Such abstraction is especially important in cases where multiple processes are simultaneously encoded in the same neural population (Caruso et al., 2018; Rigotti et al., 2013). Supervised methods like CCA (Hotelling, 1936; Safaie et al., 2023), dPCA (Kobak et al., 2016a), RRR (Izenman, 1975) do achieve abstraction since they optimise behaviour decoding rather than reconstruction. However, they were not created for time-series data and thus lack explicit dynamics modelling.

Addressing this limitation, dynamical approaches (Churchland et al., 2012; Duncker and Sahani, 2021; Ahamed et al., 2021; Mante et al., 2013; Pandarinath et al., 2018b; Durstewitz, 2017) model neural dynamics using the concept of a state space from physics (Strogatz, 2019). The goal is to identify a minimal set of variables that fully determine the system of interest’s state and its future dynamics. For instance, the oscillations of a pendulum can be adequately modelled in a two-dimensional state space defined by the position and momentum of the pendulum, and one can ignore properties irrelevant to the oscillations, such as colour, mass or ambient lighting. Current dynamical methods, in contrast, model the activity of the entire neural population, which may not be relevant to understanding a given behaviour. While valuable as in silico proxies for empirical studies (Durstewitz et al., 2023), their representations may contain too much detail to be interpretable. For example, the dynamics of different processes may become tangled (Russo et al., 2018). Behaviour-based approaches may help untangle neural representations and render them meaningful, as suggested by (Krakauer et al., 2017).

One strategy for improving interpretability is partitioning the latent space into behaviourally relevant and irrelevant subspaces, as done in subspace methods (Hurwitz et al., 2021; Sani et al., 2021, 2024; Li et al., 2024). However, in all these methods, the dynamics of the subspaces are still coupled, preventing us from modelling dynamics in the relevant subspace without reference to the irrelevant subspace.

Our method, BunDLe-Net (Behaviour and Dynamics Learning Network), explicitly learns an abstraction of neural dynamics as a state-space model restricted to specific behavioural contexts, in line with the principles of the neuroconnectionist research programme (Doerig et al., 2023). Our framework naturally emerges from applying a causal notion of consistent abstractions to dynamical systems (Grosse-Wentrup et al., 2024; Rubenstein et al., 2017). Theoretically, this abstraction allows causal statements about the neural dynamics under specific conditions. Empirically, BunDLe-Net learns trajectories that form compact bundles, which reveal branching points on the neural manifold corresponding to behavioural decisions, often preceding external behaviour.

We apply BunDLe-Net to calcium imaging data from *C. elegans* (Kato et al., 2015), electrophysiological data from the rat hippocampus (Grosmark and Buzski, 2016), and multi-unit activity from the monkey somatosensory cortex (Chowdhury et al., 2020) (see Supplementary Fig. S9), demonstrating its broad applicability across species, modalities, and behaviours. In *C. elegans*, BunDLe-Net uncovers behavioural motifs in the latent manifold which align with established behavioural patterns, e.g. the run-pirouette motif (Pierce-Shimomura et al., 1999). These manifolds show that the organism probabilistically switches between deterministic behavioural motifs at a few highly reproducible decision points. In rat hippocampal data, BunDLe-Net identifies recurrent cycles in neural activity, capturing intrinsic dynamics often missed by other methods.

BunDLe-Net is versatile across data types, handling both trial-based and freely moving animal data while modelling behaviour that is discrete, continuous, or a combination thereof. It also embeds recordings from separate sessions or animals into a shared latent space, revealing consistent manifolds across organisms. By providing a structured view of how neural activity shapes behaviour, BunDLe-Net offers a flexible framework for uncovering fundamental principles of neural computation.

## Results

### 1 Fundamental principles and architecture

Our approach to manifold learning is to construct a minimal model of behaviour from neural activity. We firstly require this model to preserve maximal information about the behaviour. However, to serve as a *model* of behaviour, it is not enough to capture behavioural information alone; the representation must also be able to predict how the system evolves, including how future behaviour unfolds from the current state. To this end, we secondly require the latent state of our model at each time point to capture all information in the neural dynamics that is relevant to predict the immediate future. We achieve this by constructing a Markovian latent embedding, such that the state at any moment is a sufficient representation of the system’s dynamics.

Together, these conditions yield a latent representation that is both self-contained and behaviourally meaningful. We implement this framework using a two-armed architecture, BunDLe-Net (Behaviour and Dynamics Learning Network), shown and explained in Fig. 1. We show in the methods section that under certain conditions the model learned by this architecture from purely observational data can be interpreted as a causal model.

**Figure 1:**
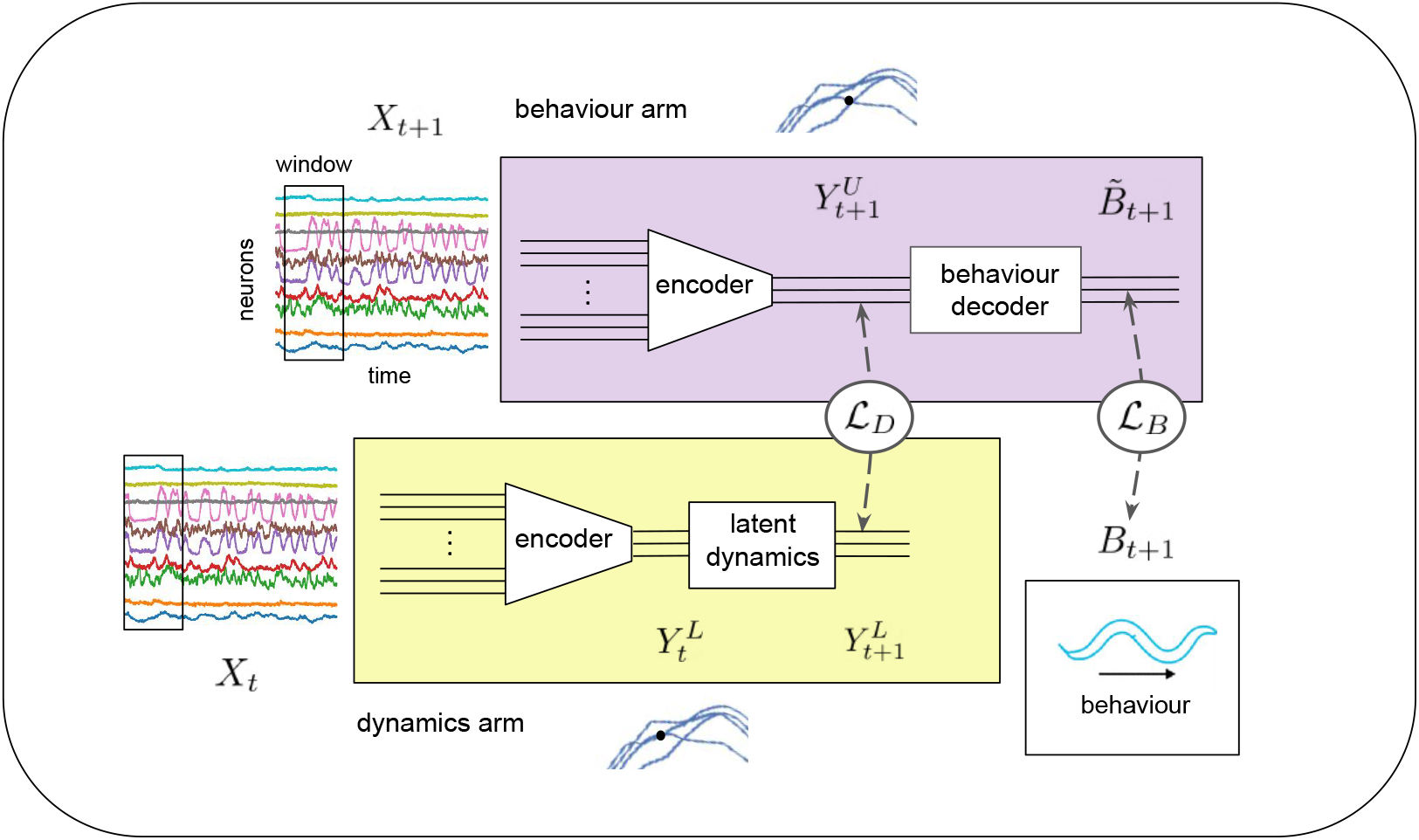
BunDLe-Net architecture. The architecture consists of the behaviour (upper) and dynamics (lower) arm. In the dynamics arm, a high-dimensional vector of neural activity (or a time window thereof) *X*_*t*_ is embedded to a point in latent space 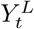 via an encoder. Then, a first-order latent dynamics model predicts the next latent state, 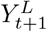, which is then compared to the true latent state, obtained by encoding the observed neural state *X*_*t*+1_ via the same encoder, yielding 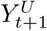 in the behaviour arm. The loss function *L*_D_ quantifies the discrepancy between the predicted and true latent states. Jointly training the encoder and the latent dynamics model to minimise ℒ_D_ results in a low-dimensional latent trajectory *Y*_*t*_ that is Markovian, meaning that the current state contains all necessary information to predict the next state, with no dependence on past states. Simply enforcing Markovianity, however, is insufficient to produce a meaningful latent representation, as trivial solutions (e.g., constant latent state / point embedding) are also Markovian. To address this, we require that our latent space preserves information corresponding to a chosen behaviour of interest. This is enforced by a loss function *L*_B_ that penalises the reconstruction error between the true behaviour (*B*_*t*+1_) and those predicted from the latent representation 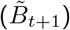. By jointly minimising ℒ_D_ and ℒ_B_, the architecture learns a Markov representation of neural dynamics specific to our behaviour of interest.

### 1.1 Dimensionality of latent space

A common assumption in current NML methods is that more latent dimensions mean more information preserved (Gao et al., 2017; Elmoznino and Bonner, 2024; Sheng et al., 2022; Vaccari et al., 2024). While this might be true for linear methods like PCA, things are more subtle for non-linear methods: mathematically, any high-dimensional space can be mapped one-to-one onto a one-dimensional space, albeit with discontinuities, as per Nettos theorem (Sagan, 1994). While such mappings preserve information, they are not necessarily a meaningful representation of the data. This raises the question of which latent dimensionality is best suited to reveal behaviourally-relevant structure in the neural dynamics.

Fortunately, there is an elegant solution to this question. In the state-space of a dynamical system, trajectories with different future outcomes are not allowed to intersect since at the intersection point, it would be impossible to predict which of the paths the system takes from the current state alone; this fact from dynamical systems theory (Strogatz, 2019) is also known in neuroscience as trajectory tangling (Russo et al., 2018). If we want to interpret a latent embedding as a faithful representation of the system’s dynamics, we thus need to embed neural activity in a space with sufficiently high dimensionality to allow trajectories to unfold without intersections. In one and two dimensions, there exist patterns of dynamics for which intersections are unavoidable (Kuratowski, 1930). However, remarkably, a result from graph theory tells us that we can embed any finite graph in three dimensions without the edges having to intersect (Cohen et al., 1995). Since transitions between a discrete set of behaviours form a finite graph, this result means that the dynamics between any set of behaviours can be embedded in three dimensions without trajectory crossings, provided that we have a sufficiently non-linear model to realise this embedding. This surprising insight has the additional benefit that three-dimensional embeddings can be visualised and are thus intuitively interpretable. We thus fix the latent space dimension in BunDLe-Net to three for visualisation purposes, while also exploring the effect of latent dimensionality on quantitative metrics.

## 2 Validation of BunDLe-Net on C. elegans data

We applied BunDLe-Net to calcium imaging and behavioural data from *C. elegans* (Kato et al., 2015) and compared it with manifolds learned by PCA (Pearson, 1901; Kato et al., 2015), LDA (Fisher, 1936; Cunningham and Yu, 2014b), t-SNE (Dimitriadis et al., 2018b; Liu and Vinck, 2022), CEBRA-hybrid (Schneider et al., 2023), (which was shown to outperform LFADS (Pandarinath et al., 2018b; Keshtkaran et al., 2022), UMAP (McInnes et al., 2018), piVAE (Zhou and Wei, 2020a), demixed-PCA (Kobak et al., 2016b), and pfLDS (Gao et al., 2016)). In addition, we evaluated other strategies including autoencoders (Kramer, 1991; Altan et al., 2021; Ran et al., 2022), dynamics autoencoder (Chen et al., 2022; Wahlström et al., 2015), and RNN autoencoder (Pandarinath et al., 2018b). This selection spans common modelling principles (linear vs. non-linear, static vs. dynamic, unsupervised vs. supervised, RNN vs. contrastive methods), allowing us to isolate the effect of how these assumptions shape the manifold’s geometry.

Since the structure of the learned manifolds are highly consistent across worms for each method (see Supplementary Fig. S2), we show in Fig. 2.a–d the learned manifolds for one representative worm plotted in three dimensions, where the colours correspond to locomotor behaviour. For reconstructive-based methods, such as PCA (Fig. 2a), t-SNE (Fig. 2b), autoencoder (Fig. 2c), and RNN autoencoder (Fig. 2d), we observe a drift in the trajectories that obscures the recurrent nature of the dynamics. Since these methods preserve maximum variance in the data, they embed the drift (which has high variance), possibly at the cost of low-variance signals that encode information relevant to behavioural dynamics. In contrast, in methods with some form of supervision, such as LDA (Fig. 2e), dynamics autoencoder (Fig. 2f), CEBRA (Fig. 2g), and BunDLe-Net (Fig. 2h), we see that this drift is largely absent, and the recurrent dynamics become apparent. Common to these methods is that they do not aim to reconstruct the entire neural activity, but only a particular characteristic in the data: behaviour for LDA, future change in neural activity for dynamics autoencoder; behaviour and timestamp in CEBRA. Though behaviour is clearly represented in the manifolds of LDA, CEBRA, and the dynamics autoencoder, the variance in the trajectories for a given behaviour is quite large, which obscures the intrinsic dynamical structure. In contrast, BunDLe-Net produces compact bundles of trajectories that are well-separated from one another. Within each bundle the variance is low, while a high variance is observed between different bundles. This bundling reveals recurrent and branching aspects of the trajectories that were indiscernible in other methods. Such recurrence facilitates reasoning about higher-order patterns in the data, such as behavioural motifs, which we discuss in Section 4.

**Figure 2:**
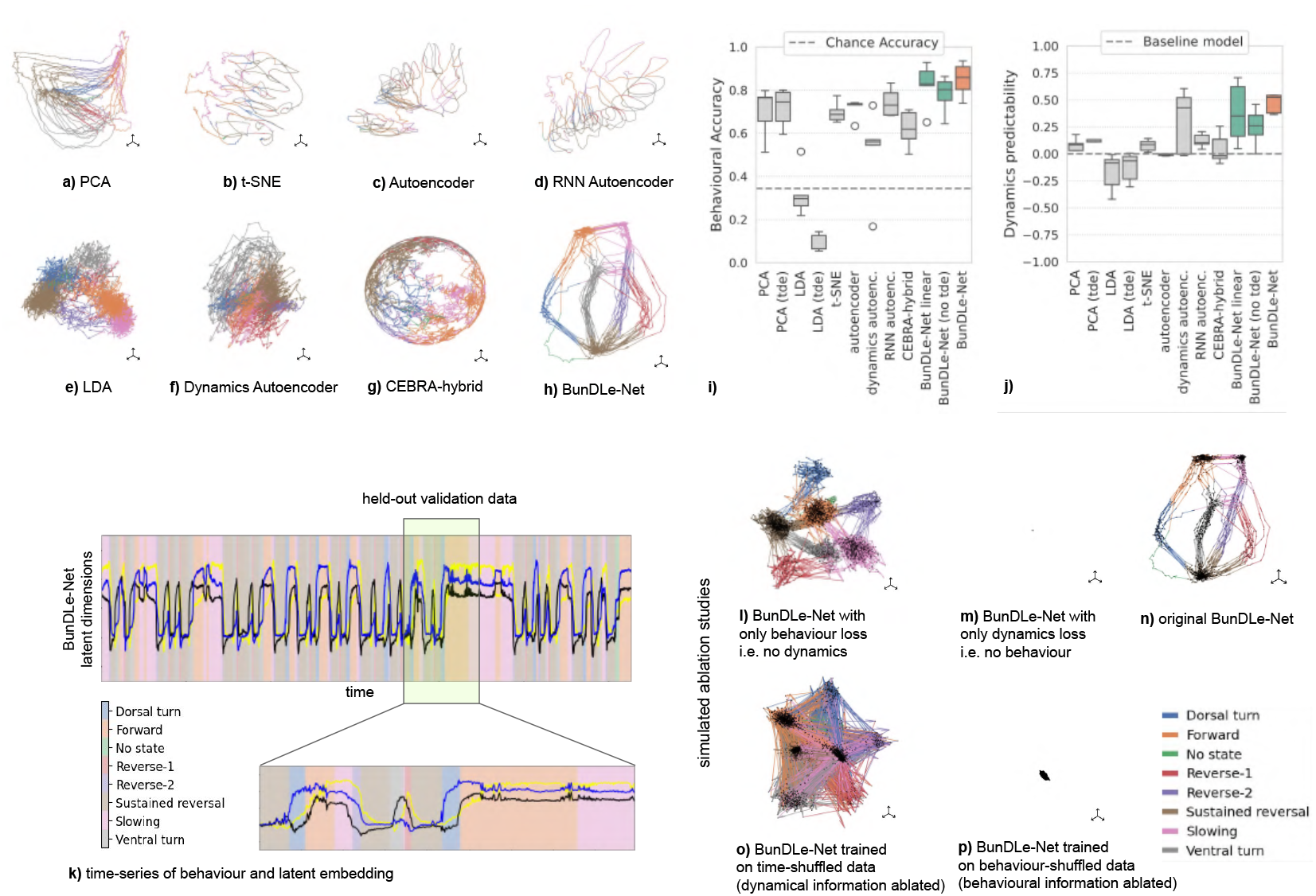
Comparison and evaluation of different manifolds on calcium imaging data from *C. elegans*. **a-h**, Neural manifolds from various NML techniques BunDLe-Net against other NML methods: **a**, PCA, **b** t-SNE, **c**, autoencoder, **d**, RNN autoencoder, **e** LDA, **f**, dynamics autoencoder, **g**, CEBRA-hybrid, and **h**, BunDLe-Net (our method). Trajectories are embedded in a 3-dimensional latent space and colour-coded according to the discrete behavioural (locomotor) states of the worm (legend at bottom right of figure). **i-j**, Quantitative evaluation of embeddings over five worms: Static methods (PCA, LDA) were augmented with time-delay embeddings that consider a past window of neural activity. BunDLe-Net was evaluated with both linear and non-linear embeddings, with and without time-delay embeddings, to assess the impact of model assumptions. **i**, Behavioural decoding accuracy from different manifold learning techniques, compared against chance-level accuracy (dotted line). **j**, Dynamics predictability on the manifold, measured via the mean squared error (MSE) of a feed-forward network predicting *Y*_*t*+1_ from *Y*_*t*_, normalised relative to a baseline model (dotted line) which simply copies the input *Y*_*t*_ to the output as the predicted value for *Y*_*t*+1_ **k**, Correspondence between the behavioural time-series and the BunDLe-Net embedding of neural activity. The plot in the foreground represents the time-series of 3-latent dimensions on BunDLe-Net’s embedding. Background shading colour indicates discrete worm behaviours. The time-series with the yellow box denotes held-out validation data, and the rest of the data served as training data. **l-p**, BunDLe-Net manifolds from simulated ablation studies on model losses (top row) and data (bottom row). Black dots represent the embedded data points. Manifold learned with **l**, only behaviour loss, **m**, only dynamics loss and **n**, both losses. **o**, Manifold trained on data with dynamical information ablated: neural and behavioural traces shuffled together to destroy temporal information. **p**, Manifold trained on data with behavioural information ablated: behavioural labels are shuffled, breaking the link between neural and behavioural data.

Importantly, this intrinsic structure of the manifold only emerges when learning a joint embedding for the neural dynamics and the behaviour, as demonstrated through model ablation studies where the BunDLe-Net model was trained with only the behaviour and dynamics loss separately (Fig. 2l-m). With a purely behavioural model (no dynamical loss), the manifold aggregates into separate clusters (Fig 2l). Each of these clusters represents information about one of the discrete behaviours. There are, however, no smooth transitions between clusters that reveal how the neural dynamics coordinate behaviours. With a purely dynamical model (no behavioural loss), the manifold collapses to a point, demonstrating that our model has the capacity for abstracting away irrelevant information (Fig. 2m). This point embedding achieves a dynamical loss of zero, since it trivially has all information about its own future at every time point. It is only when we combine both losses that we obtain the smooth transitions with clear decision points in Fig. 2n. These smooth transitions are also seen in Fig. 2k, which shows BunDLe-Net’s latent states constantly evolving towards the next behavioural state.

## 3 Branching trajectories and decisions

The BunDLe-Net manifold in Fig. 2.h shows distinct branching points in the trajectory prior to an externally observed change in behaviour. This suggests that BunDLe-Net extracts internal representations from neural activity that encode for future decisions. To further study the correspondence between branching points and presence of information, we attempt to decode the future behaviour from the latent state (Fig. 3). To this end, we normalised behavioural bouts of varying durations to lie in the time interval [0, 1], allowing trajectories from different bouts to be analysed together (Fig. 3, middle panel). From each point in this latent time-series, we then fitted a decoder to predict the subsequent behavioural state (e.g., for ventral turn 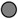 in Fig 3a, the subsequent behaviour is either slowing 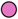 or forward 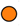). We see the cross-validated decoding accuracy increases (Fig 3, lower panel) only when the branching occurs in the latent time-series (Fig. 3 middle panel), confirming that the branching point in the manifold corresponds to when information about the future behaviour appears in the latent state representation. Notably, this result identifies when information becomes accessible in the latent state, implying that the corresponding information in the high-dimensional neural data must be present at least by this point.

**Figure 3:**
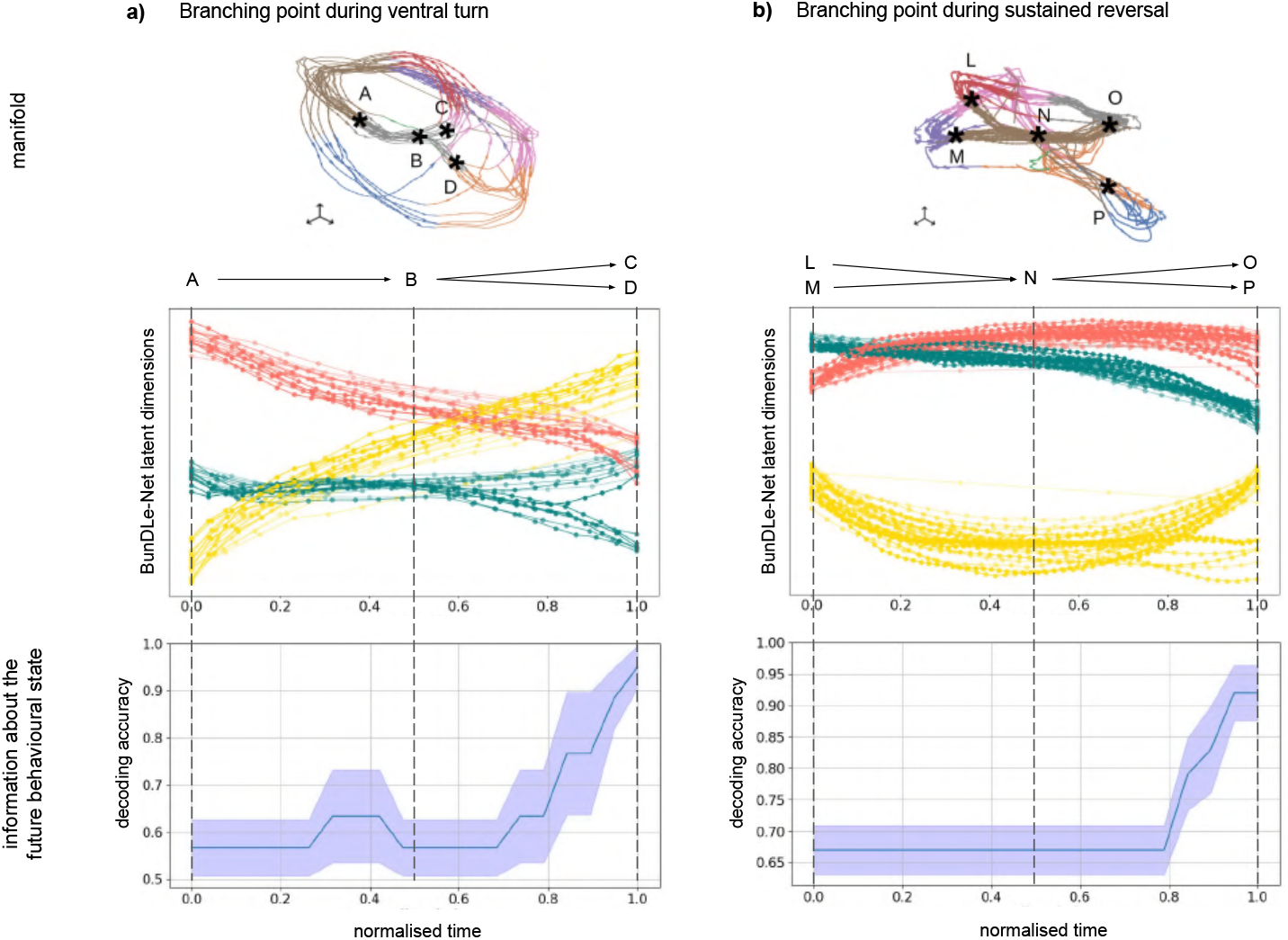
Analysis of the two branching points on the BunDLe-Net manifold that occur during sustained reversal and ventral turn. The upper panels display two views of the same manifold’s branching trajectories. The middle panels are time-series plots showing the evolution of the three latent variables during the given behaviour. To account for varying durations of each behavioural bout, we normalise time for each behaviour to the interval between 0 and 1. This enables comparison of trajectories across bouts, regardless of the speed at which their internal representations evolve. The black markers labelled (A, B, C, D and L, M, N, O, P) on the manifold correspond to the dotted lines in the time-series plot, indicating the beginning, middle, and end of the behavioural bouts. The lower panels show the cross-validated decoding accuracy for predicting the future behaviour from the manifold over normalised time. The shaded area corresponds to variation of the decoding accuracy between the cross-validation folds.

## 4 Shared latent space facilitates comparison of embeddings

Next, we show how BunDLe-Net can learn a shared latent space from data recorded from multiple organisms or sessions, while allowing for varying durations of recording in a freely-behaving organism, and different subsets of neurons recorded. By keeping the behaviour decoder and latent dynamics modules fixed across five worms, and learning only the encoder for each worm, we are able to project to a shared latent space, enabling us to compare manifolds across organisms. Examining Fig. 4a, we observe a similar branching topology in the trajectories of all five worms. Along a given branch, the dynamics are deterministic, with probabilistic decisions confined only to the branching points. Specifically, in worm-1, we identify five prominent branches (Fig. 4b) with stereotypical behavioural motifs, which we label *C*_1_, …, *C*5. Motif *C*_1_, …, *C*_4_ correspond to the well known pirouette sequence in literature (Pierce-Shimomura et al., 1999), while *C*_5_ contains the run. These behavioural motifs form the building blocks of the neural manifold and are also consistent with the ones found by (Grosse-Wentrup et al., 2024).

**Figure 4:**
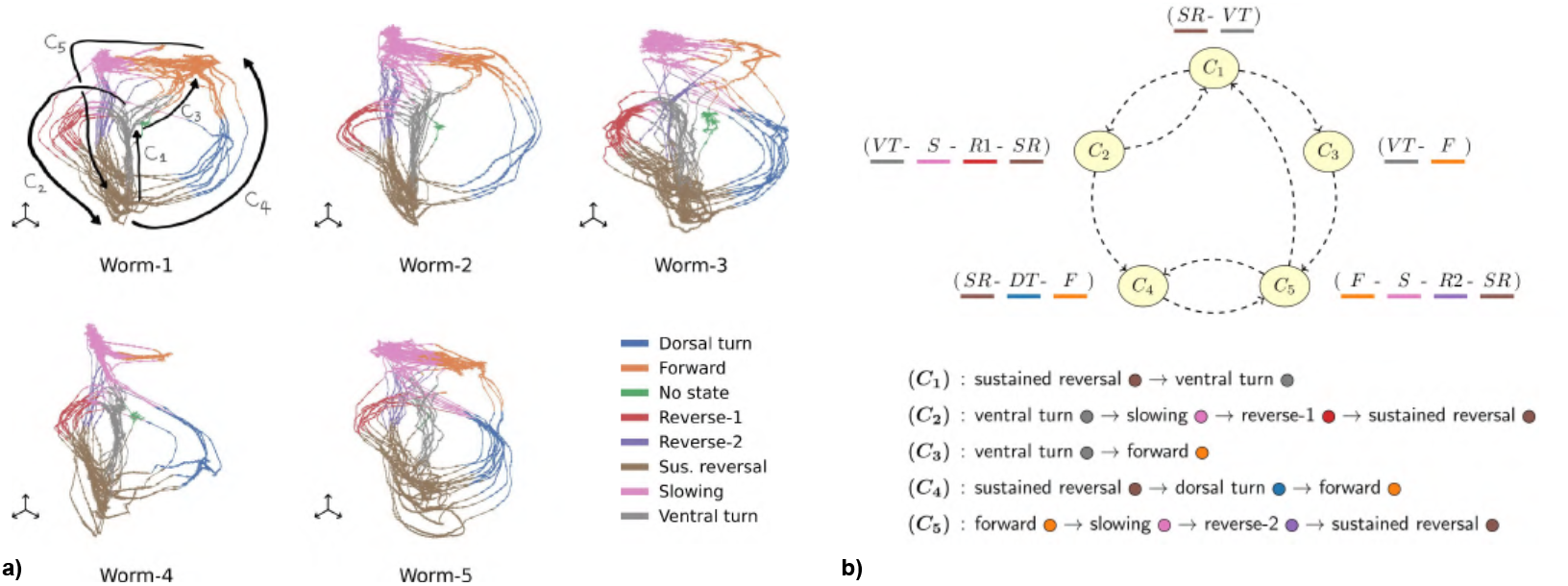
Shared latent space and analysis of branching topology in terms of behavioural motifs. **a**, BunDLe-Net manifolds of five worms in a shared latent space. The discrete locomotor behaviour is colour-coded. By keeping the behaviour decoder and latent dynamics modules fixed, we can map neural data from different worms into a common latent space, even when different sets of neurons are recorded across worms. For worm-1, the branches (segments between branching points) of the manifold are labelled *C*_1_, …, *C*_5_. Each of these branches constitutes a deterministic behavioural motif. **b**, Reduced transition diagram between behavioural motifs of worm-1. Since the manifold consists of stereotypical deterministic motifs between which the worm switches at branching points, we can reduce it to this Markov transition diagram, where each node corresponds to a branch or behavioural motif. This exploits the fact that stochasticity is confined to the branching points in the BunDLe-Net manifold.

The shared latent space further shows that the behavioural motifs are largely consistent across worms (Fig. 4b). For example, motif ***C***_**2**_ consistently forms a loop 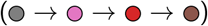 that is present in all worms. Motifs ***C***_**1**_ and ***C***_**5**_ are also consistently present across worms. However, motif ***C***_**4**_ is absent in Worm-4, where both Worm-4 and Worm-5 instead exhibit a different motif: sustained reversal 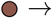 dorsal turn 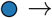 slowing 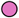. These variations may result from recording times being too short to capture all possible transitions for a given animal or varying individual strategies.

Since the probabilistic transitions are locally confined on the manifold, we can further abstract our manifold to a transition diagram between motifs. In this diagram, the transitions are purely probabilistic since we have captured as much possible deterministic behaviour within a motif. This transition diagram summarises the neural manifold’s structure and also aligns with findings from purely behavioural studies (Costa et al., 2024; Stephens et al., 2011).

## 5 Decoding multiple behaviours from rat data

To demonstrate the flexibility of BunDLe-Net across different modalities of neural and behavioural data, we applied our method to a publicly available rat electrophysiology dataset that has been widely as a benchmark for latent methods (Zhou and Wei, 2020b; Schneider et al., 2023). The dataset consists of CA1 neural and behavioural recordings (position and direction) from rats navigating a 1.6 m linear track (Grosmark and Buzski, 2016). Since the rats deterministically traverse the track in alternating directions, the behavioural manifold consists of two parallel linear trajectories, one per direction, connected at the track ends, thus forming a loop. The neural manifold, if it encodes this behaviour and keeps only behaviourally-relevant information, should have a similar structure. The known structure of the ground truth behavioural manifold allows us to validate whether BunDLe-Net correctly recovers this topology.

The same BunDLe-Net architecture in Fig. 1 was used with minimal modifications to accommodate the new behavioural paradigm (see Methods). Fig. 5a shows the neural manifold of four rats by BunDLe-Net and other algorithms. We see that manifolds from supervised methods, such as RRR, CCA, CEBRA-hybrid and BunDLe-Net, show a higher correspondence with the behaviour, as can be seen from the coloured embeddings where similar behaviours are localised near each other. The unsupervised methods, which prioritise full-state reconstruction, show less structure in this sense. Focussing on the embeddings of rat-1 (Achilles), we see that only CEBRA and BunDLe-Net recover the closed loop structure as expected by the behavioural task of the animal. In contrast to BunDLe-Net’s embedding of *C. elegans* data (Fig. 4a), there are no branching points in the trajectory of the rat’s neural dynamics, due to the deterministic nature of the task. The simplicity of the task also results in a manifold topology that can be embedded in just two dimensions. Although we provided all methods with three latent dimensions, only BunDLe-Net collapses the representation onto a two-dimensional subspace, while the other methods fail to identify the lower-dimensionality of the manifold.

**Figure 5:**
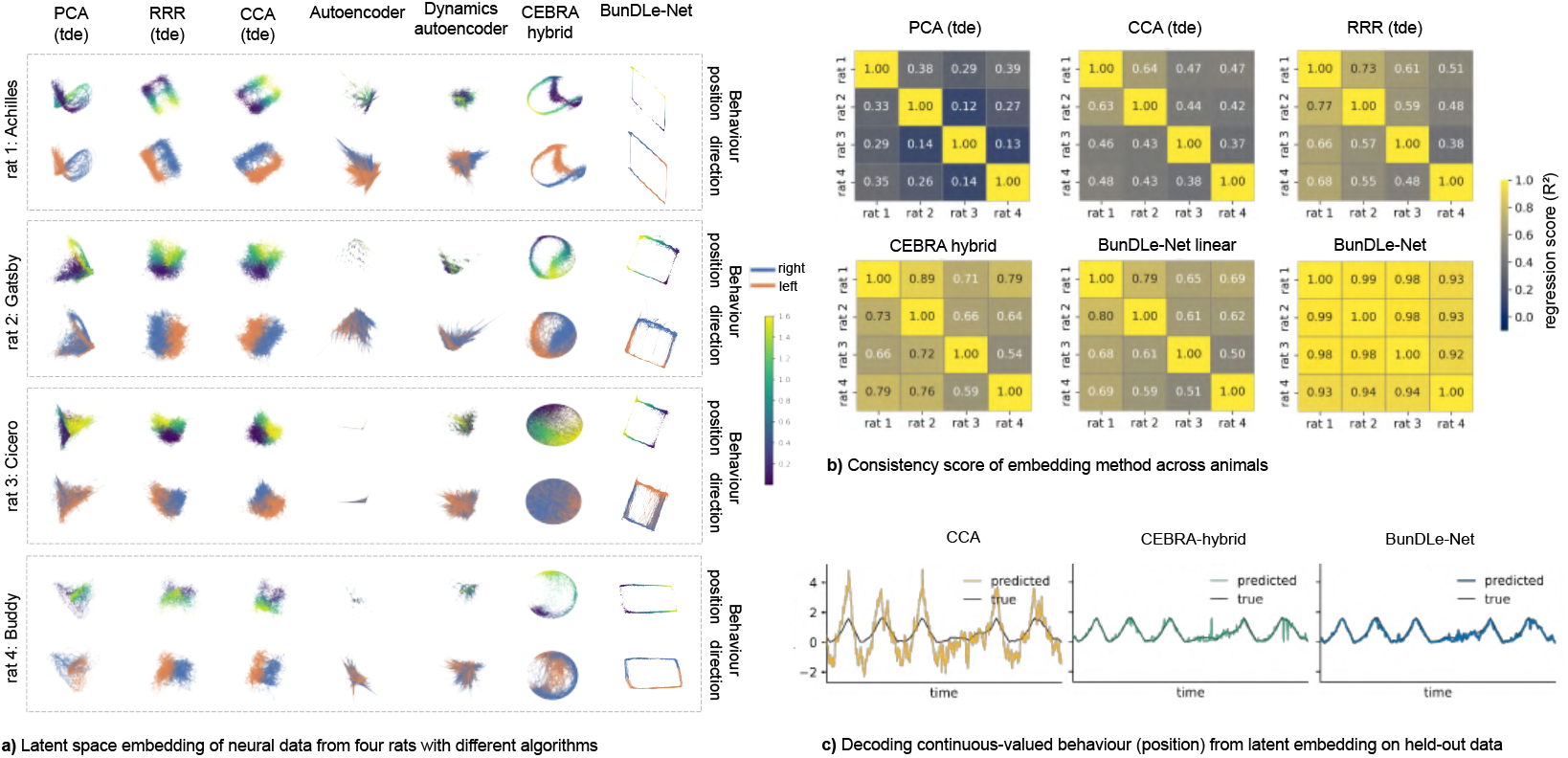
**a**, Comparison of neural manifolds from rat hippocampal electrophysiology and behavioural recordings. Data from four rats are shown in separate grey dashed boxes, with columns corresponding to different embedding methods. To enable fair comparison with neural network-based approaches, linear methods (PCA, RRR and CCA) are augmented with a time-delay embedding (tde) that incorporates temporal context. For each method, the same embedding is shown twice, coloured by position (continuous colour map) and movement direction (binary colour map), respectively, illustrating how behaviour is represented in the learned latent space. The top row shows embedded data points, while the bottom row connects successive time points to visualise how the trajectories evolve over time. **b**, Consistency analysis across animals. Matrices show pairwise linear regression scores (*R*^2^) between behaviour-aligned latent embeddings from different rats. For each method, latent trajectories were aligned to behaviour by binning the continuous behavioural variable and averaging embeddings within each bin. Behaviour-matched embeddings from each animal pair were then compared using ordinary least-squares regression. Higher *R*^2^ values indicate greater cross-animal consistency of the learned latent geometry. **c**, Decoding behaviour from latent embeddings. We compare a linear model (CCA), a non-linear model (CEBRA), and BunDLe-Net on held-out test data. Each method embeds previously unseen neural activity into its latent space, and behaviour is then decoded directly from the resulting embedding. The true behavioural trace is shown in black, and the coloured traces show the decoded behaviour for each method.

Turning our attention to the other rats, we see that BunDLe-Net’s embedding remains consistent in terms of topological structure, i.e., a rectangle where one dimension encodes position, and the orthogonal one encodes direction. In contrast, the other embeddings become obscured by noise, resulting in a barely discernible topological structure. BunDLe-Net’s trajectories also show smoother temporal transitions, whereas other embeddings exhibit discontinuous jumps that obscure dynamic structure. This contrast may be attributed to BunDLe-Net’s architecture, which is designed to preserve dynamical information.

To quantify the consistency of our manifolds across animals, we measured how well the behaviour-aligned manifold learned in one rat predicts the manifold of another (see Methods). As shown in Fig. 5b, supervised linear approaches (CCA and RRR) outperform the unsupervised baseline (PCA). A linear variant of BunDLe-Net outperforms other linear methods, indicating that the performance of our method is not solely due to nonlinearity. Incorporating nonlinearity further improves performance, with BunDLe-Net showing the best consistency across animals. In Fig. 5c, we assess the manifolds based on how well they predict the behaviour (rat’s position) from unseen neural data. Our method achieves high decoding performance, consistent with the well-established role of hippocampal activity in encoding spatial position (Grosmark and Buzski, 2016). While decoding is not the primary objective of our approach, these results provide validation that the learned representations preserve behaviourally relevant information while remaining smooth, noise-robust, and consistent across animals.

## Discussion

Neural manifolds are typically evaluated in terms of compression and decoding, i.e., how well they represent high-dimensional neural activity and how accurately they predict behaviour. Yet neural data analysis has long aimed for the more ambitious goals of understanding how distributed neural activity realises computations and causes behaviour (Vyas et al., 2020; Langdon et al., 2023; Jazayeri and Afraz, 2017; Perich et al., 2025). In most existing approaches, manifold geometry is not explicitly tied to questions of causation, and when causality is considered at all, it is often conceived as a static, structural property between neurons in local circuits, assessed via focal stimulation or lesion experiments (Houweling and Brecht, 2008; Otchy et al., 2015). This has led to a tension between the dynamic and distributed nature of neural manifolds and the static, local view of causal influence.

Our work offers a complementary perspective: causally interesting structure is not uniformly distributed along the manifold but is concentrated at specific points in time and space, namely, the branching points at which trajectories diverge toward different future behaviours. The rest of the manifold, while dynamically important, primarily serves to coordinate the evolution of the system between such decision points. By learning Markovian embeddings that abstract away dynamics irrelevant to a given behavioural context, BunDLe-Net uncovers a manifold geometry in which most trajectories form low-variance bundles corresponding to deterministic behavioural motifs, punctuated by reproducible branching points where future behaviour becomes probabilistic. This organisation is particularly clear in the *C. elegans* data, where we find stereotyped motifs that align with known locomotor patterns, and in the rat hippocampus, where simple loop trajectories align with task structure.

This view reframes how causality can be investigated in population activity. Rather than asking which individual neurons cause a behaviour, our results suggest that causally effective perturbations should be understood as state- and time-dependent interventions on coordinated patterns of activity. Because BunDLe-Net provides a low-dimensional, Markovian summary of the behaviourally relevant dynamics, it allows us to identify when information about alternative futures first becomes available in the neural state. Our experimental results suggest that at these branching points, small perturbations to the appropriate population activity are likely to alter the subsequent trajectory of the system and thereby change behaviour, while the same perturbation may have little or no effect when applied away from these points. Concretely, BunDLe-Net enables the derivation of testable causal predictions on the effects of spatio-temporal interventions on neural activity, where the predicted effect on neural dynamics and behaviour depends on the global neural perturbation and on the position of the neural dynamics on the manifold at the time of intervention. To test these predictions, one could design closed-loop stimulation protocols that target specific branching points on the manifold to steer the neural state to bring about a desired behaviour. This strategy moves beyond classical stimulation paradigms and aligns with work emphasising global, distributed encoding of behaviour (Rigotti et al., 2013; Caruso et al., 2018), in which no single neuron or small subset is sufficient on its own.

From a computational standpoint, the manifold perspective also offers a bridge between neural dynamics and algorithms. If we think of a computation as a sequence of internal decisions that transform inputs into outputs (Vyas et al., 2020; Sussillo and Barak, 2013), then the branching structure of the manifold can be interpreted as a coarse description of an algorithm implemented by the neural system: deterministic motifs correspond to stereotyped computational subroutines (Driscoll et al., 2024), and branching points correspond to conditional operations where alternative processing paths are selected. Different tasks, strategies, or learning histories should then give rise to different branching topologies and motif repertoires, even if overall decoding performance is similar. BunDLe-Net thus provides a candidate tool for identifying neural algorithms not merely from the decoding accuracy but based on the structure of the latent trajectory and branching points.

While our present work focuses on the relationship between neural activity and behaviour, the framework naturally extends to include sensory inputs, internal variables, or task parameters as additional nodes in the causal graph. This would enable a more complete characterisation of how neural activity mediates the processing of stimuli to behavioural decisions. In summary, by combining principles from causal abstraction with behaviourally-relevant dynamical modelling, BunDLe-Net moves neural manifold learning beyond compression and decoding toward a structural, temporally resolved account of when and where causally relevant decisions are encoded in population activity.

## Data and code availability

The results for this paper can be found here https://github.com/akshey-kumar/bundlenet_results. An open-source Python implementation can be found here https://github.com/neuroinformatics-uni-vienna/NC-MCM/

## Methods

### 1 From theoretical principles to model architecture

Our goal here is to construct a minimal model of behaviour that can be interpreted as a self-contained state-space model. We adopt the notion of a model from the framework of Structural Causal Models (SCMs) (Pearl, 2009; Grosse-Wentrup et al., 2024), and apply it to the setting of dynamical systems. We chose this framework because it allows us to describe a system in terms of interventions, enabling us to reason about the system as a compact real-world model in its own right.

We define a dynamical SCM as follows: Let *X*_*t*_ ∈ *X* be a set of variables representing the state of the system at time *t*. We consider a collection of such sets *{X*_*t*_ : *t* ∈ N*}*, one for each time point, and associate with each *X*_*t*_ a corresponding set of exogenous noise variables *ϵ*_*t*_, with one noise variable for each component of *X*_*t*_. The variables *X*_*t*_ are endogenous, meaning they can be observed and intervened on, while *ϵ*_*t*_ are exogenous, representing unobserved sources of variation that contribute to uncertainty in the observed data.

#### Definition 1.

*(Dynamical SCM) A dynamical structural causal model (dSCM) is a tuple* 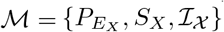 *where*,

- 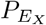 *is the joint probability distribution over the exogenous variables such that ϵ*_*t*_ *and* 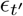 *are independent for all t t*^*′*^.
- *S*_*X*_ *is a set of update equations, one for each time point, of the form, X*_*t*_ = *f*_*t*_[*Pa*(*X*_*t*_), *ϵ*_*t*_], *where the parent set refers to all direct causes of X*_*t*_ *and is restricted to lie in the temporal past:* 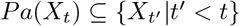.
- *I*_*X*_ *is a set of allowed interventions on the endogenous variables. An intervention on X*_*t*_ *is denoted do*(*X*_*t*_ = *x*).

The essence is that in a *dSCM*, the direct causes of a variable *X*_*t*_ are restricted to lie in its temporal past. In this framework, we allow all possible *complete* interventions, meaning interventions on all components of *X*_*t*_ jointly; formally *I*_*X*_ = {*do*(*X*_*t*_ = *x*) : *x*∈ *X}*. Note that the distribution over the exogenous variables, together with the structural equations, induce a distribution over the endogenous variables *P* (*X*_*t*_ | *Pa*(*X*_*t*_)) which we shall refer to in our definition of main criteria.

#### Neural abstractions

The full neural state dynamics can be modelled through a dSCM which we denote *M*_*X*_, where the variables *{X*_*t*_ : *t* ∈ N*}* represent the high-dimensional neural state evolving via update equations *X*_*t*_ = *f*_*t*_(*Pa*(*X*_*t*_), *ϵ*_*t*_). Rather than modelling the full neural state, our goal is to extract a lower-dimensional embedding *Y*_*t*_ = *τ* (*X*_*t*_) that is self-contained; i.e. it should evolve according to its own model *M*_*Y*_ (with its own update rules, interventions, and exogenous distribution), without reference to the original high-dimensional state *X*_*t*_. Inspired by the ideas and notation of Rubenstein et al. (2017), we identify notions of dynamical and behavioural consistency, which form the basis of our theoretical contribution.

##### 1.1 Consistency criteria

###### Definition 2.

*(Dynamical consistency) Let* 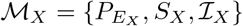 *and* 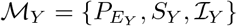 *be two dSCMs and let τ* : *X* → *Y be a surjective mapping between the state spaces of M*_*X*_ *and M*_*Y*_. *We say that the triple {M*_*X*_, *M*_*Y*_, *τ} shows observational dynamical consistency (DC) iff for all t we have, Y*_*t*_ = *τ* (*X*_*t*_), *and for all x*^*′*^ ∈ *X, y* ∈ *Y we have*^1^,

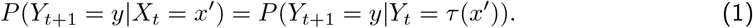

This condition can be understood as the conditional independence statement (*Y*_*t*+1_ ⊥ *X*_*t*_ | *Y*_*t*_). Intuitively, it guarantees that once we know the latent state *Y*_*t*_, the high-dimensional neural state *X*_*t*_ carries no further predictive information about the latent dynamics. In other words, the latent process evolves on its own, without reference to the original high-dimensional state. While many embeddings may satisfy dynamically consistency, not all of them are necessarily meaningful to gain insights about behaviour. This motivates a second notion, behavioural consistency, which requires the latent embedding to preserve information about behaviour. Let 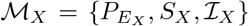 be a dSCM and let (*B*_*t*_ ∈ *B*) be a set of behaviour variables. *S*_*B*_ is a set of equations *B*_*t*_ = *g*_*t*_(*X*_*t*_, *ϵ*_*B*_) for every *t*. We define 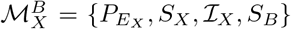 as the corresponding *dSCM in a behavioural context*.

###### Definition 3.

*(Behavioural consistency) Let* 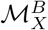 *and* 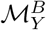 *be two dSCMs in the same behavioural context. The triple* 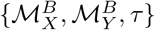 *is said to show observational behavioural consistency (BC) iff for all t we have Y*_*t*_ = *τ* (*X*_*t*_) *and for all b* ∈ *B, x* ∈ *X we have*,

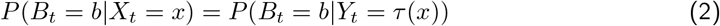

This definition ensures that both the high-dimensional and low-dimensional models contain the same amount of information about the behaviour.

##### 1.2 Linking observational and causal consistency

The above definitions specify criteria on the conditional probability distributions that can be derived from purely observational data. However, for the derived model *M*_*Y*_ to be a valid real-world causal model, i.e. to be meaningful beyond statistical associations, it must remain consistent under interventions (out-of-distribution experimental manipulation). We say that the triple {*M*_*X*_, *M*_*Y*_, *M τ}* shows *causal dynamical consistency* if the distribution under interventions respect the mapping *τ* in the following sense:

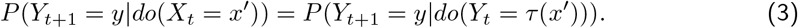

Typically, we need experiments to validate such causal statements. However, we can identify conditions under which observational statements (learned purely from data), imply the causal statement. In particular, we will show that if the underlying dSCM for neural activity, *M*_*X*_ is Markov (i.e. the only direct cause of the *X*_*t*+1_ is *X*_*t*_; *Pa*(*X*_*t*+1_) = {*X*_*t*_}), then observational dynamical consistency implies causal dynamical consistency.

##### 1.3 Conditions for causal consistency

The following lemma and theorem establish how we may learn causal dynamical consistent transformations from purely observational data in a Markov setting.

###### Lemma 1.

*Let M*_*X*_ *be a dSCM whose dynamics are Markov, i*.*e. Pa*(*X*_*t*+1_) = *{X*_*t*_*}, and let τ* : *X* → *Y be a surjective mapping. Let M*_*Y*_ *be a dSCM that describes the dynamics of Y*_*t*_ := *τ* (*X*_*t*_). *If {M*_*X*_, *M* _*Y*_, *M τ} shows observational dynamical consistency, then M*_*Y*_ *is also a Markov dSCM*.

*Proof*. ^2^ The theorem states that a sufficient condition for *M*_*Y*_ to be a Markov dSCM is *observational dynamical consistency*, i.e for all *x* ∈ *X, y*^*′*^ ∈ *Y, t* ∈ *T* ⊆ N we have,

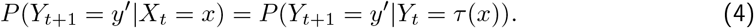

To prove that *M*_*Y*_ is Markov, we will show that the term

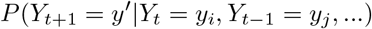

equals to

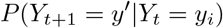

under the assumption of observational DC. First, we note the equivalence between the statements *Y*_*t*_ = *y* and *X*_*t*_ ∈ *τ*^−1^(*y*) and use this to expand,

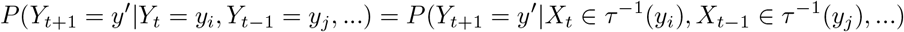

Using Bayes rule, the additivity rule for mutually exclusive events, and again Bayes rule, we express the term as a sum over past trajectories,

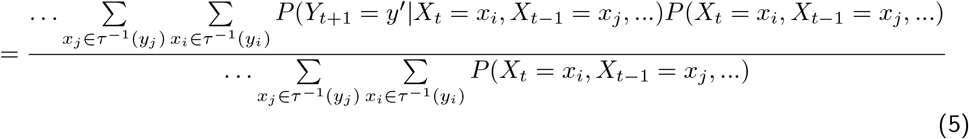

Now consider the term, *P* (*Y*_*t*+1_ = *y*^*′*^| *X*_*t*_ = *x*_*i*_, *X*_*t*−1_ = *x*_*j*_, …). Since *M*_*X*_ is Markov, it can be simplified as follows,

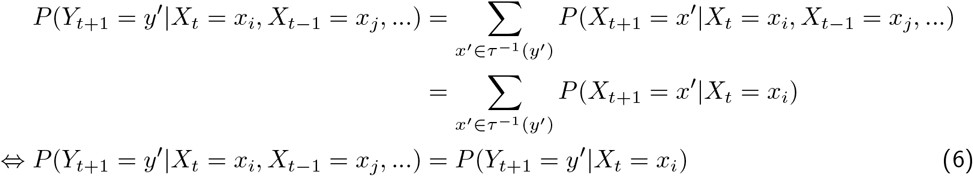

Using Equation (6), the term (5) becomes,

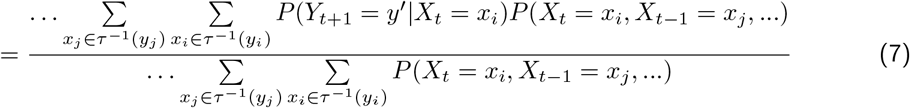

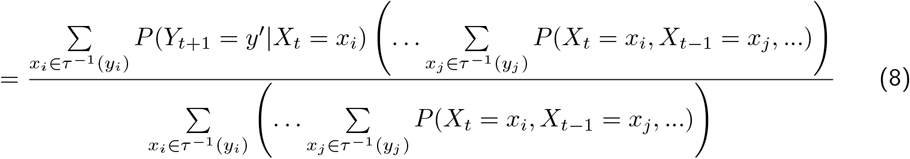

Now, if we assume that observational DC holds, for a given value of *Y*_*t*_ = *y*_*i*_ the distribution *P* (*Y*_*t*+1_ = *y*^*′*^|*X*_*t*_ = *x*_*i*_) is the same constant as long as *x*_*i*_ ∈ *τ*^−1^(*y*_*i*_). It only depends on the value of *y*_*i*_, and is thus given by *P* (*Y*_*t*+1_ = *y*^*′*^|*Y*_*t*_ = *y*_*i*_). Since the first summation sums over *x*_*i*_ for a fixed value of *y*_*i*_, we have,

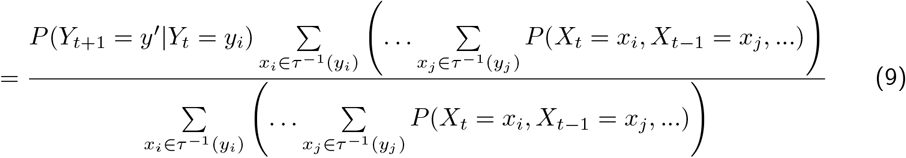

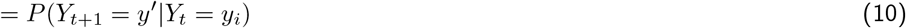

Thus, we started with the term *P* (*Y*_*t*+1_ = *y*^*′*^|*Y*_*t*_ = *y*_*i*_, *Y*_*t*−1_ = *y*_*j*_, …), used observational DC to show that it simplifies to *P* (*Y*_*t*+1_ = *y*^*′*^|*Y*_*t*_ = *y*_*i*_) if *M*_*X*_ is Markov, which means that observational DC is a sufficient condition for *M*_*Y*_ to be Markov.

###### Theorem 1.

*Let M*_*X*_ *and M*_*Y*_ *be two dSCMs such that Y*_*t*_ = *τ* (*X*_*t*_) *where τ is a surjective mapping. A sufficient condition for causal dynamical consistency of the triple {M*_*X*_, *M*_*Y*_, *τ}, is for M*_*X*_ *to be a Markov dSCM and for observational dynamical consistency to hold*.

*Proof*. Since *M*_*X*_ is a Markov dSCM and *{M*_*X*_, *M*_*Y*_, *τ}* satisfies observational DC, it follows from Lemma (1) that *M*_*Y*_ is also a Markov dSCM. We proceed by showing that the *RHS* of Equations (1) and (3) are equal if *M*_*X*_ and *M*_*Y*_ are Markov. Then we show that the *LHS* of these equations are also equal, concluding our proof.

Since *M*_*Y*_ is a Markov dSCM, we have *Pa*(*Y*_*t*+1_) = {*Y*_*t*_*}*. Since *Y*_*t*_ is the only parent of *Y*_*t*+1_ and intervening on all parents leaves the conditional distribution unchanged, we have,

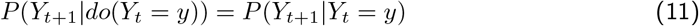

Similarly, since *M*_*X*_ is also a Markov dSCM, we have.

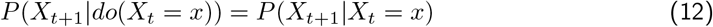

Now consider the LHS of equations (1) and (3). We denote the pre-image of *y* ∈ *Y* with respect to the mapping *τ* as the set *τ*^−1^(*y*) = *{x* ∈ *X*|*τ* (*x*) = *y}*. Consequently for any choice of *y* ∈ *Y*,

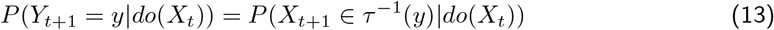

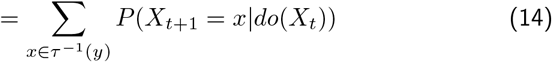

The summation follows since all the states *x* ∈ *τ*^−1^(*y*) ⊆ *X* are mutually exclusive. In a similar way we also have for any *y* ∈ *Y*,

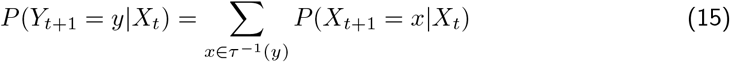

Since the summands in *RHS* of equations (14) and (15) are equal by equation (12), it follows that,

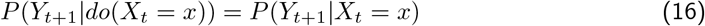

Thus, from equations (12) and (16), we see that the observational DC is sufficient for causal DC in our prescribed framework of a Markov model *M*_*X*_.

Lemma 1 shows that observational dynamic consistency implies that the Y-level model is Markov if the X-level is Markov. This is a statement about statistical dependencies, not about causal mechanisms. Theorem 1 strengthens this result by showing that, under the assumption that neural dynamics are Markovian, observational dynamic consistency also implies causal dynamic consistency. As a result, under these conditions, we can identify a consistent, causally interpretable model in the latent space directly from observational data, without needing interventions. This model retains the same causal structure as the neural-level model, with transitions governed by the Markov property.

##### 1.4 From consistency criteria to architecture: Learning transformations from data

The theoretical criteria developed so far tell us what properties a transformation *τ* must satisfy in order to yield a valid low-dimensional model. In particular, Theorem 1 shows that if the original system has Markov dynamics, a transformation satisfying observational consistency also preserves causal structure. We now turn to the practical question: how can we learn such a transformation from data? The key insight is that dynamical consistency can be expressed as a commutativity condition in a simple diagram, and this, in turn, leads directly to a learnable architecture. By enforcing commutativity during training, we can ensure that the learned transformation satisfies DC, and thus inherits the desirable theoretical guarantees.

#### Dynamical consistency in practice

Here we show that a transformation *τ* satisfies observational dynamical consistency if the diagram in Fig. 6 commutes: it does not make a difference if we take the path from *X*_*t*_ to *Y*_*t*+1_ via *X*_*t*+1_ or via *Y*_*t*_. The following theorem formalises this. **Theorem 2**. *If the diagram in Fig. 6 commutes, observational dynamical consistency is fulfilled*.

**Figure 6:**
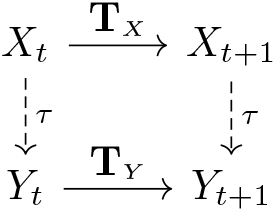
Commutativity diagram

*Proof*. For this proof, consider a probability vector *P*_*X*_ ∈ [0, 1]^|*X* |^ which represents the probability mass function at time *t* over all possible states in space *X*. We are now interested in how this probability vector can be transformed to give us a probability mass function *P*_*Y*_ ∈ [0, 1]^|*Y*|^ over the states in *Y* at the time *t* + 1. If the diagram commutes, then we may either perform the dynamics on the *X*-level and then *τ* transform, or first *τ* transform and then perform the dynamics on the considerably lower-dimensional *Y*-level.

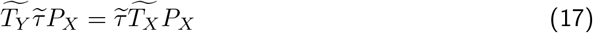

where 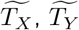 and 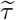 denote matrix representations^3^ of the transition and embedding models in the probability spaces of states. We define them as follows,

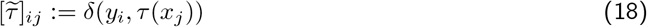

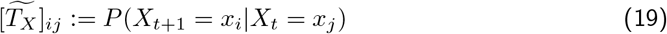

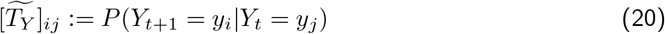

Note that fixing 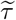 and 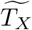 induces a transition matrix 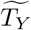 on the Y-level, which is given by 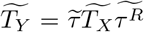, where 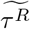 is the right inverse of non-square matrix 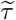. For the remainder of the proof, however, it suffices to consider the form in Equation 20.

Since Equation 17 must hold for any possible choice of the initial probability vector *P*_*X*_, we have,

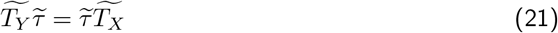

We can simplify this as follows (using the laws of matrix multiplication, the definitions and probability rules),

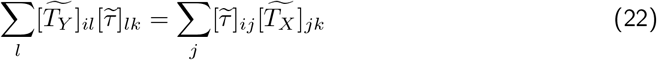

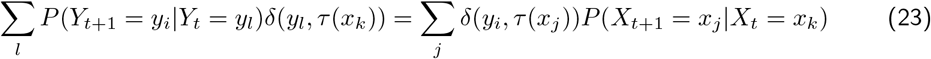

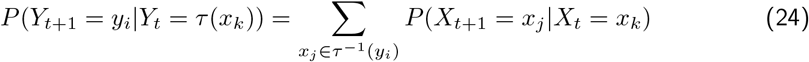

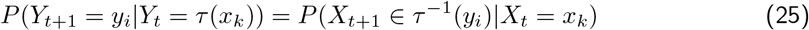

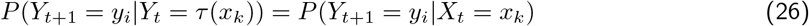

Hence, the diagram commuting implies observational dynamical consistency.

#### BunDLe-Net architecture

The preceding theorems provide the foundation for our learning architecture. They show that we can either evolve the system forward in the high-dimensional space and then apply *τ*, or apply *τ* first and then evolve forward in the low-dimensional space. We implement this commutativity using a two-armed neural network shown in Fig. 1. Since we are given time series data, we do not need to learn the high-dimensional transition map *T*_*X*_, which reduces computational cost. We need only learn *τ* and *T*_*Y*_, which correspond to the *encoder* and *latent dynamics* module in Fig. 1. In the upper arm, we map the next observation *X*_*t*+1_ into the latent space, giving 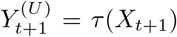. In the lower arm, we embed the current observation *Y*_*t*_ = *τ* (*X*_*t*_) and then evolve it using a low-dimensional transition model *T*_*Y*_, yielding 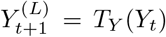. To enforce dynamical consistency, we introduce a dynamical loss term as,

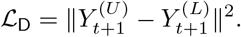

If this term converges, then by Theorem 2 the learned encoder *τ* satisfies observational dynamical consistency, and if the original system is Markovian, Theorem 1 also guarantees causal consistency. Thus, the latent representation defines a valid state-space and causal model.

#### Behavioural consistency in practice

Behavioural consistency (equation 2) requires the latent representation to preserve all information needed for predicting behaviour. To enforce this, we jointly train a *behavioural decoder* module (Fig. 1) on the latent space and add a behavioural loss term *L*_B_ alongside the dynamical loss. The estimated *Y* ^*U*^ is passed through a predictor that outputs *B*_*t*+1_. For discrete behavioural labels, we use the cross-entropy loss

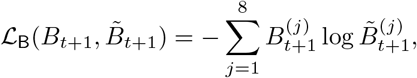

where 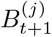 is the one-hot encoded label and 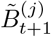 the softmax output.

The *encoder, latent dynamics*, and *behaviour decoder* modules are trained jointly to minimise the combined loss *L* = (1 − *γ*) *L* _D_ + *γ L*_B_, where *γ* is a hyperparameter between 0 and 1. Together, these terms ensure that the BunDLe-Net extracts the relevant behavioural dynamics from neural data. An open-source Python implementation is available at https://github.com/neuroinformatics-uni-vienna/NC-MCM.

#### Adaptation for continuous-valued multidimensional behaviour

For continuous-valued behavioural variables such as the rat dataset, we modify only the *behaviour predictor* module in Fig. 1 to match the dimensionality of the continuous-valued behavioural outputs. In this setting, we use a mean squared error loss for *L*_B_. All other architectural components and training parameters remain unchanged.

## 2 Worm data and preprocessing

We used whole-brain calcium imaging data from *C. elegans* as collected and preprocessed by Kato et al. (2015). This dataset is well-suited for evaluating BunDLe-Net as it captures rich behavioural variability across multiple animals. It comprises time-series recordings from five worms, each spanning 18 minutes (2.9 Hz sampling rate), yielding 2500 to 3500 time points per worm and recordings from 100 to 200 neurons. Not all recorded neurons could be identified; hence, only a subset is labelled for each worm, with different yet overlapping subsets identified across worms. Human-annotated motor behaviours were assigned at each time point, with eight discrete behavioural states: forward, slowing, dorsal turn, ventral turn, no-state, sustained reversal, reversal-1, and reversal-2. The data contains repeated transitions between states, enabling robust assessment of behavioural dynamics and flexibility.

Motor behaviours in the dataset were annotated based on the activity of a small set of well-characterised neurons (AVAR, AVAL, SMDVR, SMDVL, SMDDR, SMDDL, RIBR, RIBL), whose firing patterns reliably correspond to specific motor states even in immobilised worms (Kato et al., 2015). To prevent circularly inferring behaviour from neurons used to define it, we excluded these neurons from all analyses. We treat them as a ground-truth behavioural readout rather than as part of the input population.

The data was temporally filtered with a low-pass filter of cut-off frequency of 0.05 Hz to mitigate high-frequency noise. To take into account the history of neural activity, we allowed the input to consist of a sliding window of past neural activity rather than a single time point. Window length was treated as a tuneable hyperparameter.

## 3 Learning modules

### Architecture parameters

The *encoder* layer *τ* of our network consists of a series of dense ReLU layers (Agarap, 2018) consisting of 50, 30, 25, 10 units, a linear layer of 3 units, followed by a normalisation layer. For the *predictor* and *T*_*Y*_ layer, we use a single dense linear layer each. This sufficed to achieve good performance on the datasets used in this work. For *T*_*Y*_, we also add a normalisation layer so that the output remains in the scaled in the latent space learned by the encoder.

### Gaussian noise against overfitting

To safeguard against overfitting of the model, we introduce Gaussian white noise in the latent space by incorporating it in the encoder layer *τ*. Injecting Gaussian white noise is a well-established regularisation technique that makes the model robust to overfitting (Camuto et al., 2020; Zur et al., 2009). Since we are working with relatively limited data in the context of artificial neural networks, guarding against overfitting becomes particularly important.

### Model validation

To determine the optimal parameters for the model, including the number and types of layers, we use a held-out validation set on worm-1. The neural and behavioural data of worm-1 is partitioned into seven folds along the time axis, and one fold is randomly selected as the validation set from the time-ordered dataset. The remaining data forms the training set. By choosing an entire fold in the data as a validation set, we ensure that the model performs as well on unseen data. This would not be the case if we created our validation set by *iid* (independent and identically distributed) sampling due to high time correlations in the time series. After selecting the optimal model parameters through validation on worm-1, we train models with the same parameters on the other worms. Since we only use worm-1 for parameter tuning, and the model performs well on other worms as seen in Fig. 4 and Fig. S2, we can be confident that the chosen parameters generalise well.

### Training details

Training was performed with the ADAM optimiser (Kingma and Ba, 2014) with a learning rate of 0.001 and batch size of 100. The *γ* parameter of BunDLe-Net was chosen to be 0.9 to ensure that *L*_D_ and *L*_B_ are roughly the same order of magnitude during training. We trained BunDLe-Net until the losses converged.

## 4 Quantitative evaluation metrics

We evaluate a latent space representation based on how well it preserves behavioural and dynamical information. To estimate the *behavioural information* of an embedding, we train a simple feed-forward neural network in a supervised setting to predict behaviour from the embedding. We use a network consisting of a single linear layer since it already demonstrated a high decoding accuracy (~ 0.94) on the raw neural traces. The decoding accuracy from the latent state is used as a metric for the information content about *B* in the embedding, with the decoding accuracy obtained on the raw, high-dimensional neural traces serving as the baseline. To evaluate the *dynamical information* in the embedding, we train an autoregressor network to predict *Y*_*t*+1_ from *Y*_*t*_. The mean squared error between the predicted and true *Y*_*t*+1_ is estimated. From this, we compute a predictability metric for the dynamics, defined as 1 − MSE_*m*_*/*MSE_*io*_, where MSE_*m*_ is the mean squared error of the model and MSE_*io*_ is the mean squared error of a trivial autoregressor that copies its input to the output (baseline performance). For each worm, we trained the evaluation models on a training set of the embedded data and performed the evaluation on a held-out test set to prevent overfitting. This train test split is separate from the model validation procedure used for selecting the BunDLe-Net’s parameters described in 3.

## 5 Comparable embeddings across worms

To produce comparable embeddings, we first trained a model on each worm separately. We then extracted the latent dynamics layer *T*_*Y*_ and behaviour predictor layer from the model with the least loss (worm-1, in this case). We then trained fresh models on each worm, with the chosen *T*_*Y*_ and behaviour predictor layers from worm-1 frozen in, until the losses converged. Thus, the new models would only have to learn the encoder *τ* for each worm while the other layers remained unchanged throughout the learning process. Note that this approach was feasible despite recording different neurons from each worm. Adopting this strategy allows us to find consistent embeddings across the worms, should they exist. The embeddings are illustrated in Fig. 4.

Note that instead of projecting the neural data from worms 2 to 5 into the latent space of worm 1, we can also simply fit separate models on each worm’s data, which we do in Supplementary Fig. S2. Due to differing initialisations of BunDLe-Net this results in visually different embeddings. Though visually different, these embeddings share the same topology, which is seen to be independent of the initialisation. For ease of visual comparison between embeddings, we adopt the shared latent space approach to produce comparable embeddings.

## 6 Comparison with other algorithms

Our goal is to understand how modelling assumptions affect the topology of the manifold and which properties they induce. To this end, we decomposed existing approaches into their fundamental components, including full state reconstruction, dynamics reconstruction, non-linear encoding, and behavioural supervision, and examined these components in isolation. This allows us to identify which modelling elements most strongly influence the embedding without conflating their effects.

We selected a range of methods that collectively span the building blocks of current techniques (see Supplementary Fig. S1). These include standard unsupervised approaches like PCA and t-SNE; a supervised linear method, LDA; and deep learning models such as an autoencoder (reconstruction through a bottleneck) and a dynamics autoencoder (dynamics reconstruction through a bottleneck). Importantly, all deep learning models were given access to a tuneable temporal window of past neural activity, rather than just a single time point, allowing them to leverage temporal structure where relevant. We also include CEBRA variants, which use contrastive learning and have outperformed multiple recent methods, including LFADS, AutoLFADS, UMAP, and pi-VAE (which itself outperforms LFADS, demixed PCA, and PfLDS) (Schneider et al., 2023; Zhou and Wei, 2020a).

### 6.1 Hyperparameter tuning and Model selection

We tuned all parametrised models using Bayesian optimisation as implemented in Ray Tune (Liaw et al., 2018), which is a standard framework for hyperparameter search. Each model was optimised independently with respect to its own performance metric, using 200 trials drawn from a search space informed by literature. We used the optimal hyperparameters to instantiate and train each model. To account for variability due to random initialisation, we trained five randomly initialised models and selected the one with the lowest loss. This ensures that we obtain the best-performing model. Below is a list of the hyperparameter search spaces and metrics used.

#### t-SNE

For t-distributed stochastic neighbour embedding (t-SNE), we tuned parameters including perplexity (5 to 50), early exaggeration (12 to 32), and the number of optimisation iterations (250 to 2,500) and the temporal window parameter between 1 (single time slice) and 20 (a 20-frame window). The optimal configuration was selected based on the Kullback-Leibler (KL) divergence between the high-dimensional and embedded distributions, which is the native objective minimised by the algorithm.

#### CEBRA

For CEBRA, we tuned parameters related to both model architecture and training dynamics. Specifically, we tuned over predefined configurations, including the default, provided in the CEBRA documentation: offset10-model and offset1-model. Other hyperparameters included batch size (256 or 512), learning rate (log-uniformly sampled between 10^−5^ and 10^−3^), temperature for the contrastive loss (uniformly sampled between 0.1 and 1.0), and the number of training iterations (ranging from 2,000 to 10,000). We also explored different time offset values (from 5 to 20 timesteps) and distance metrics (cosine or Euclidean). Model quality was assessed using a behavioural decoding task, as described in the original CEBRA study: a k-nearest neighbours classifier was trained on the learned embedding to predict behavioural labels, and test-set classification accuracy was used as the selection criterion. After determining the optimal hyperparameters, five models were trained, and the best model was chosen based on the loss.

#### Autoencoder

For the autoencoder, we considered a variety of encoder architectures with varying depth and width (Table 1), and the decoder architecture was always set as the mirror of the encoder architecture. The bottleneck of the autoencoder was fixed to 3 dimensions in order to facilitate comparisons across models. We tuned learning rate (from 10^−4^ to 10^−1^), number of training epochs (log-uniformly sampled from 10 to 500), batch size (log-uniformly sampled from 32 to 512), and the temporal window parameter (log-uniformly sampled from 1 to 20). The performance metric was the mean squared error (MSE) between the input and its reconstruction, reflecting the fidelity of the reconstruction.

**Table 1:** Encoder architectures where each entry lists the number of units per layer from input to bottleneck. Note that the number of units in the bottleneck is fixed to 3 to facilitate comparison.

| Architecture | Layer Sizes |
| --- | --- |
| Shallow | 50, 10 |
| Increasing then Decreasing | 100, 150, 50, 10 |
| BunDLe-Net encoder | 50, 30, 25, 10 |
| Deep | 100, 80, 60, 40, 20, 10, 10 |
| Intermediate | 75, 25, 10 |

#### Dynamics autoencoder

The dynamics autoencoder models the dynamics by learning to predict the change in neural activity Δ*x*_*t*+1_ = *x*_*t*+1_ − *x*_*t*_ given the neural activity at time *t*, using a bottleneck architecture. We used the same search space and architectural variants as for the autoencoder. Model performance was again assessed by the MSE between predicted and true activity at time *t* + 1.

## 7 Analysis of branching points on the manifold

To relate branching points in the latent trajectories to decisions about future behaviour, we performed a time-resolved decoding analysis on behaviour-specific latent trajectories. We focused on behaviours for which a clear branching structure was observed in the learned manifold, in particular, ventral turn and sustained reversal. From the behavioural annotations, we extracted all contiguous bouts of the selected behaviour from the training data, together with the corresponding latent trajectories. Behavioural bouts varied in duration, so to enable comparisons across bouts, we normalised time within each bout to the interval [0, 1] and interpolated the latent trajectories to a fixed length of 20 time steps. Interpolation was performed independently for each latent dimension using piecewise linear interpolation.

To quantify how information about future behaviour evolves along the latent trajectory, we decoded the behaviour immediately following each bout from the latent state at each interpolated time point. Specifically, for each normalised time point, we trained a linear classifier (logistic regression) to predict the subsequent behavioural state from the latent representation. Decoding performance was evaluated using 5-fold cross-validation, and accuracy was computed independently at each time point along the trajectory. The resulting time-resolved decoding accuracy reflects how information about the upcoming behaviour becomes accessible in the latent state as the trajectory approaches and passes the branching point.

## 8 BunDLe-Net on rat hippocampal data

We use a publicly available rat electrophysiology dataset from a study conducted Grosmark and Buzski (2016). This dataset has been widely employed to assess different neural embedding techniques (Schneider et al., 2023; Zhou and Wei, 2020b). It consists of simultaneously recorded electrophysiological and behavioural data as the rat navigates a 1.6-meter linear track. It is markedly different from the *C. elegans* dataset since we have spiking neural data (discrete-valued) instead of calcium image data (continuous-valued). Also, there are two behaviour variables that are simultaneously recorded and correspond to the rat’s position and running direction (left or right). The position variable is continuous-valued, while the direction is discrete-valued. Another significant difference is that the neural recordings are only from a small subset of the brain. Hence, this data set is more challenging with respect to finding a latent embedding that is causally sufficient for the neural dynamics.

The BunDLe-Net architecture required minimal modifications to accommodate this new behavioural paradigm. Since we have two behaviour variables, one of which is continuous, we used a linear layer with two outputs along with a mean-squared error loss.

### 8.1 Hyperparameter tuning of algorithms on rat data

We compared the manifolds of several algorithms including principal component analysis (PCA) (Pearson, 1901), canonical correlation analysis (CCA) (Hotelling, 1936; Safaie et al., 2023), reduced rank regression (RRR) (Izenman, 1975), CEBRA (Schneider et al., 2023), autoencoder (Kramer, 1991; Altan et al., 2021; Ran et al., 2022), and dynamics autoencoder (Chen et al., 2022; Wahlström et al., 2015). The linear methods (PCA, CCA and RRR) do not require hyperparameter tuning and were applied using their standard formulations. For all other methods, hyperparameters were optimised on the rat dataset as described below.

#### CEBRA

The rat hippocampus dataset used in this study was also a primary benchmark in the original CEBRA work. We therefore used the hyperparameter configuration reported in the original publication, which had already been optimised on this dataset. Specifically, we used the *offset10* model architecture with a three-dimensional embedding. Training was performed using a batch size of 512, learning rate of 3 *×* 10^−4^, temperature parameter set to 1, and a cosine distance metric. The model was trained for a maximum of 5,000 iterations using time-conditional contrastive learning with a temporal offset of 10. Hybrid training was enabled, which means that the model considers both behavioural and temporal information.

#### Autoencoder

For the autoencoder baseline, hyperparameter tuning was performed using Bayesian optimisation implemented in Ray Tune. We considered a variety of encoder architectures with varying depth and width (Table 1), and the decoder architecture was always set as the mirror of the encoder architecture. The bottleneck of the autoencoder was fixed to 3 dimensions in order to facilitate comparisons across models.

The hyperparameter search space included the learning rate, sampled log-uniformly from [10^−4^, 10^−1^]; the number of training epochs, sampled log-uniformly from [10, 1000]; the batch size, sampled from a log-uniform distribution between 32 and 512; and the length of the temporal input window, sampled log-uniformly from [1, 20] time steps. Models were trained to minimise the mean squared reconstruction error, which served as the optimisation objective. Bayesian optimisation was run for 200 trials, and the configuration achieving the lowest validation mean squared error was selected.

#### Dynamics autoencoder

Hyperparameter tuning for the dynamics autoencoder followed the same optimisation procedure and search space as for the standard autoencoder.

### 8.2 Behaviour alignment and consistency score

For the rat embeddings, we evaluated whether each method yielded behaviourally consistent latent representations across animals by comparing embeddings across all pairs of rats. For each animal and method, latent trajectories were aligned to the continuous behavioural variable by binning behaviour into 1600 evenly spaced bins and averaging latent vectors within each bin. Behaviour-aligned embeddings from two rats were then matched on shared behavioural bins, yielding pairs of latent vectors corresponding to the same behavioural states. To quantify cross-animal consistency, we fit an ordinary least-squares linear regression mapping the behaviour-aligned embedding of animal *i* to that of animal *j* and report the coefficient of determination (*R*^2^). A high *R*^2^ indicates that the geometry of the embedding in one animal spans a linear subspace sufficient to reconstruct the other animal’s embedding. Because linear predictability is a relatively stringent criterion, strong *R*^2^ values provide compelling evidence that the method captures shared, generalisable structure between the manifolds. The resulting matrices of regression scores therefore provide a quantitative measure of how consistently a given method represents neural activity across animals.

## Acknowledgements

We would like to thank Sebastian Tschiatschek and Simon Rittel for enriching discussions at the Causal Representation Workshop 2021, which was hosted at the Faculty of Computer Science, University of Vienna. We would also like to thank Manuel Zimmer and his lab, especially Kerem Uzel, for collaborating with us and providing the neural calcium imaging data from *C. elegans*. We also thank Verity Cook for discussions about the illustrations and figures.

This work was supported under the CHIST-ERA grant (CHIST-ERA-19-XAI-002), by the Austrian Science Fund (FWF) (grant reference I 5211-N) and the Engineering and Physical Sciences Research Council United Kingdom (grant reference EP/V055720/1), as part of the Causal Explanations in Reinforcement Learning (CausalXRL) project.

This work has been funded by the Vienna Science and Technology Fund (WWTF) [Grant ID: 10.47379/LS23070]

## Supplementary Information

### S1 Consistent latent model for linear dynamics

Here, we demonstrate how one can construct a low-dimensional dSCM for a system with linear dynamics. Note that this is not a unique solution, and there may be other abstractions that fulfil our requirements. We use the theoretical devices developed in the previous sections to present an analytical solution for the linear case and prove how it satisfies our criteria. The resulting latent model preserves the Markov causal structure and is sealed off from the full dynamics since it preserves sufficient information about its own dynamics.

### S1.1 Analytic solution for linear models

Here, we present a transformation of a linear dynamical system in a behavioural context and prove that it is both DCC and BC. Let 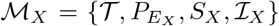 be a Markov dSCM where 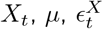 are vectors in ℝ^*n*^ and *M*, Σ are *n × n* matrices:

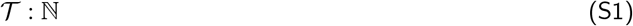

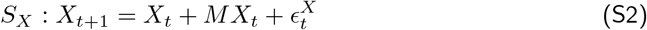

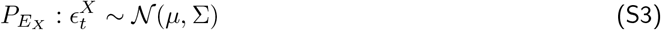

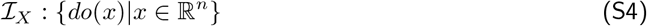

*M* represents the adjacency matrix of directed connections between various *X*^*i*^ i.e. *M*_*ij*_ ≠ 0 iff there is a causal connection, 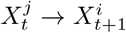. We do not allow any cycles in graph defined by *M*. We also define a behaviour based on the microvariable state as follows,

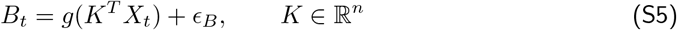

where *g* can be any arbitrary functional mapping and *ϵ*_*B*_ is a noise variable. Note that the behaviour depends on *X*_*t*_ only through the linear combination *K*^*T*^ *X*.

Now let *τ* : ℝ^*n*^→ *F* be a surjective coarse-graining^4^ to ⊆ *F* ℝ^*m*^, a lower-dimensional space. We choose *m* to be equal to the length of the longest path in the DAG defined by *M*. We represent *τ* as an *m × n* matrix such that *Y*_*t*_ := *τ* (*X*_*t*_).

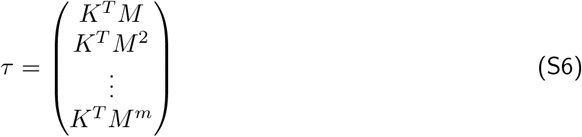

Let 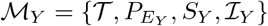 be a Markov dSCM

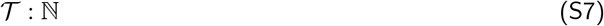

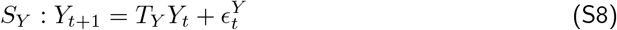

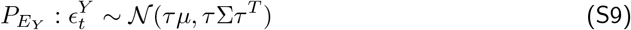

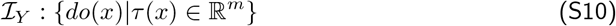

where *T*_*Y*_ is an *m × m* matrix

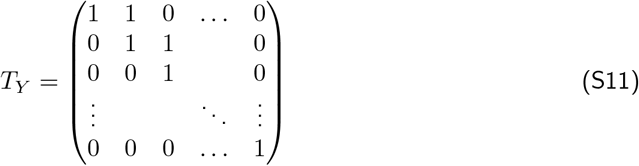

*Proof*. We begin by showing that the diagram in Theorem 2 commutes, thus ensuring observational dynamical consistency. We then invoke Theorem 1 to prove causal dynamical consistency.

#### Upper arm of the commutativity diagram

First, we apply the transition model as stipulated by *M*_*X*_ to *X*_*t*_, followed by coarse-graining with *τ* to obtain,

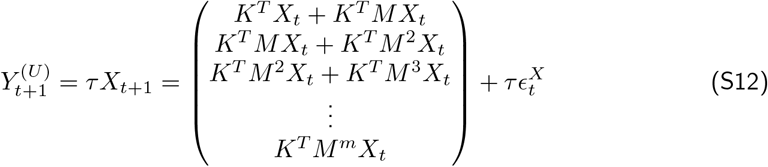

Note that the final entry in the column vector simplifies to *K*^*T*^ *M* ^*m*^*X*_*t*_ since *M* ^*m*+1^ = 0. This is because *M* is the adjacency matrix of an acyclic graph and *m* is the length of the longest possible walk in the graph.

#### Lower arm of the commutativity diagram

For this path, we first coarse-grain *X*_*t*_ with *τ* followed by applying the transition model on the *Y*-level as stipulated by *M*_*Y*_ to obtain,

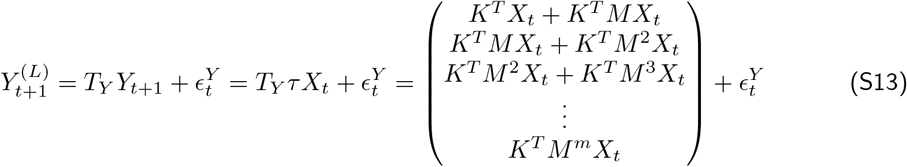

By definition of 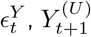 and 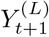 are identically distributed, thus concluding our proof of DOC. Since *M*_*X*_ is a Markov dSCM by definition, we invoke Theorem 1 to prove DCC. Behaviour causal consistency is ensured since the first component of *Y*_*t*_ is *K*_*T*_ *X*_*t*_, which contains all information about *B*_*t*_ in *X*_*t*_.

For an arbitrary coarse-graining of variables at the X-level, it may not be possible to have a description of the variables on the Y-level in terms of a dSCM. In some cases, a dSCM description might be possible, but only with a different causal structure or with latent confounders. With the above example, we have clearly shown how a dSCM can be abstracted to the Y-level in such a way that causal structure and behavioural information are preserved.

### S2 Relation of BunDLe-Net to existing methods

We present BunDLe-Net as a generic architecture in the sense that each of its modules (the encoder, latent dynamics model, and behaviour decoder) can be realised by any model of choice whether linear or non-linear, realised via neural networks or other machine learning techniques. In this paper, we have used a neural network implementation with a non-linear encoder, a linear latent dynamics model, and a linear behaviour decoder. We chose the simplest possible components that were necessary to produce a good fit to our data and generalise on held-out data. More complex model choices for the encoder and decoder, such as RNNs, might provide incremental improvements at the cost of interpretability and risk of overfitting. In general, one must adapt the modules of BunDLe-Net according to the use case and data paradigm.

In order to show how various existing manifold learning techniques are related, we consider the modifications required to convert BunDLe-Net into another manifold learning technique. This resulting Fig.S1 illustrates both how individual methods arise from BunDLe-Net and how they relate to one another through shared or inherited assumptions.

**Figure S1:**
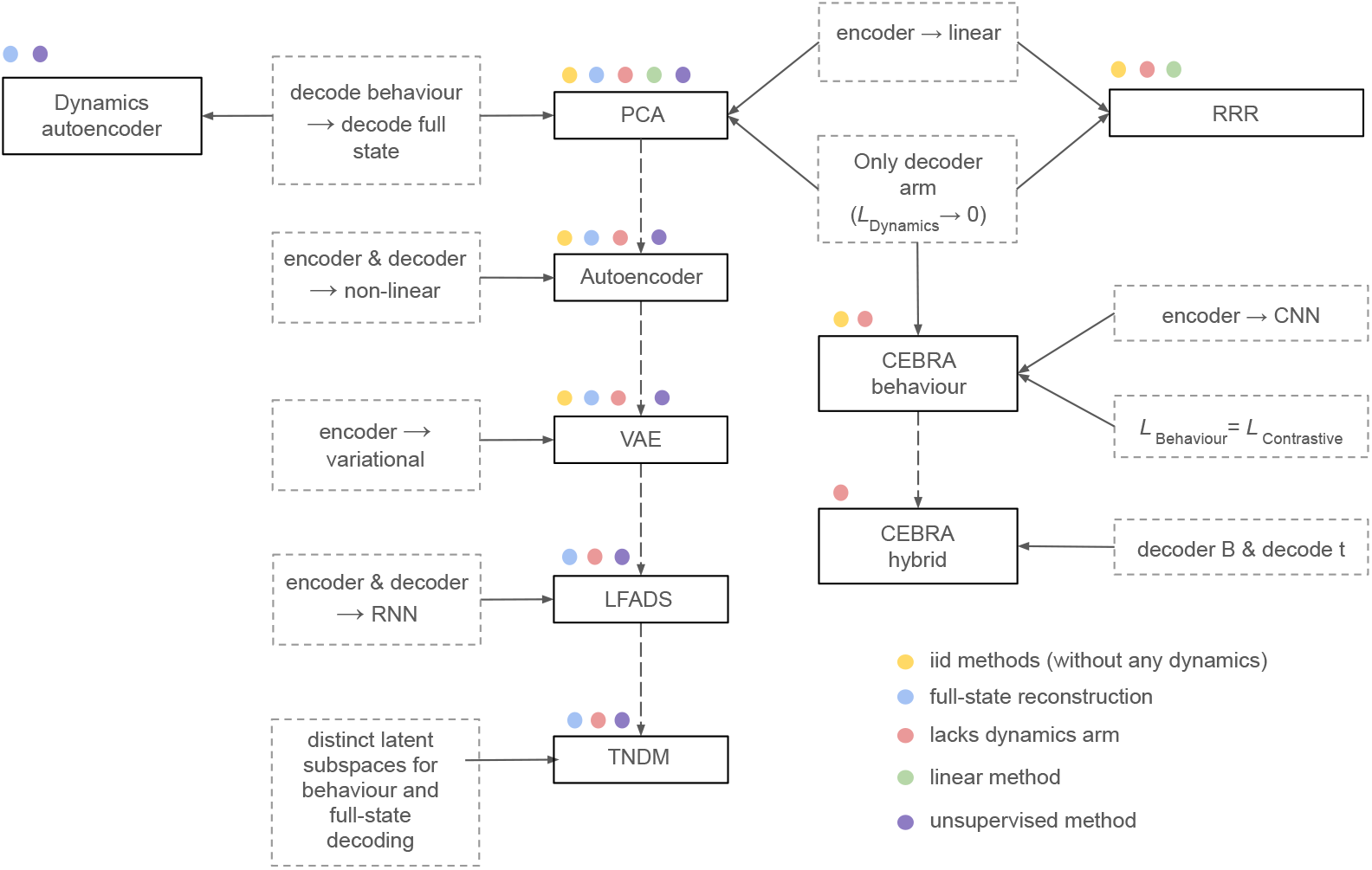
Schematic of the modifications necessary to convert our instantiation of BunDLe-Net to other commonly-used manifold learning methods. The methods are represented by solid black boxes, while the required modifications are shown in the dotted grey boxes. A solid arrow between a modification and method denotes that this is one of the modifications needed to make BunDLe-Net equivalent to a specific method. A dotted arrow between two methods indicates that the latter method inherits all modifications of the preceding method. The solid arrows have precedence over the dotted arrows in cases where the assumptions conflict.

For example, consider reduced rank regression (RRR) (Izenman,1975). This can be obtained from BunDLe-Net, by using a linear encoder and removing the dynamics loss while retaining the linear behaviour decoder. If instead, we start with BunDLe-Net, replace the behaviour decoder by a full state decoder, remove the dynamics loss, and use only linear layers, the resulting architecture becomes equivalent to principal component analysis (PCA) (Pearson, 1901). Taking PCA, and allowing the encoder and decoder to be non-linear, yields a model equivalent to a standard autoencoder (Kramer, 1991).

More generative dynamical approaches can be obtained by further modifying the architecture. Using a variational encoder on a standard autoencoder yields a variational autoencoder (VAE) (Zhou and Wei, 2020b; Li et al., 2024). Now, if we augment the VAE with an RNN encoder and decoder, and replace the sliding temporal window (for time delay embedding in BunDLe-Net) with experimental trials, we would obtain LFADS (Pandarinath et al., 2018b). If we then modify LFADS to include behaviour decoding with a separate latent subspace for behaviour and full-state decoding, we would get TNDM (Hurwitz et al., 2021).

CEBRA models (Schneider et al., 2023) differ from BunDLe-Net by three modifications. It does not have a dynamics loss that enforces Markovianity in the BunDLe-Net embedding, uses a contrastive loss based on behaviour and additionally used a convolutional neural net (CNN) as the encoder. The variant CEBRA hybrid included both behaviour and time labels in the contrastive loss.

Thus far, we have only considered models that lack the dynamics arm in Fig. 1. In our comparisons and baselines, we include another model that includes the dynamics arm but does not perform behaviour decoding. It takes as input the neural state *X*_*t*_ but instead of decoding the behaviou,r it decodes the full state at the next time step, *X*_*t*+1_. We refer to this model as a *dynamics autoencoder* and note related formulations in prior work (Chen et al., 2022; Wahlström et al., 2015). Thus, this schematic provides us an overview of existing methods, and how they are related to each other and BunDLe-Net by their modelling assumptions. Based on the modelling and data requirements, we can use this schematic to choose an appropriate instantiation of our architecture.

### S3 C. elegans manifolds on all worms

**Figure S2:**
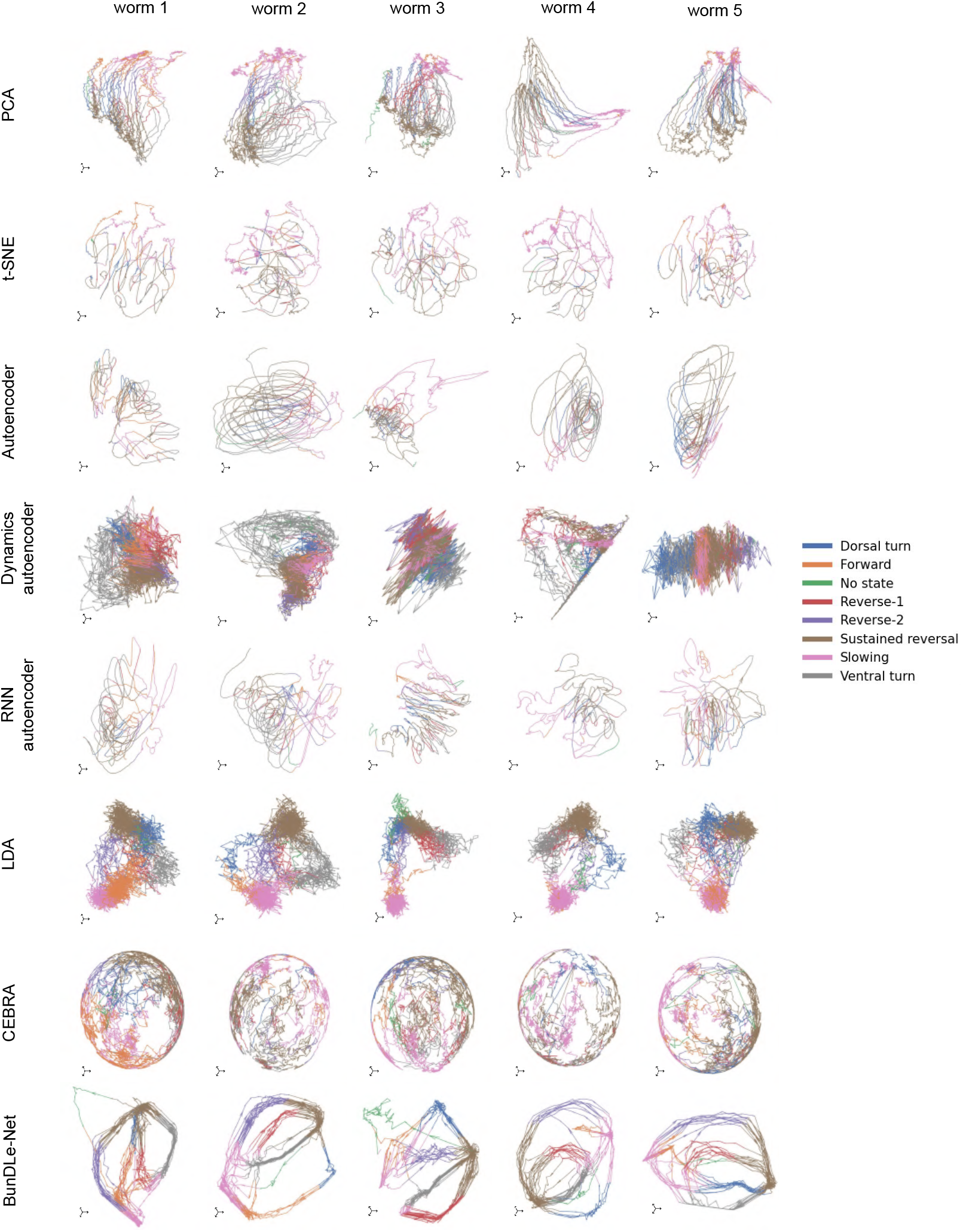
Embeddings in three dimensions of all five *C. elegans* worms for a variety of algorithms.

### S4 Effect of varying the BunDLe-Net parameter gamma

**Figure S3:**
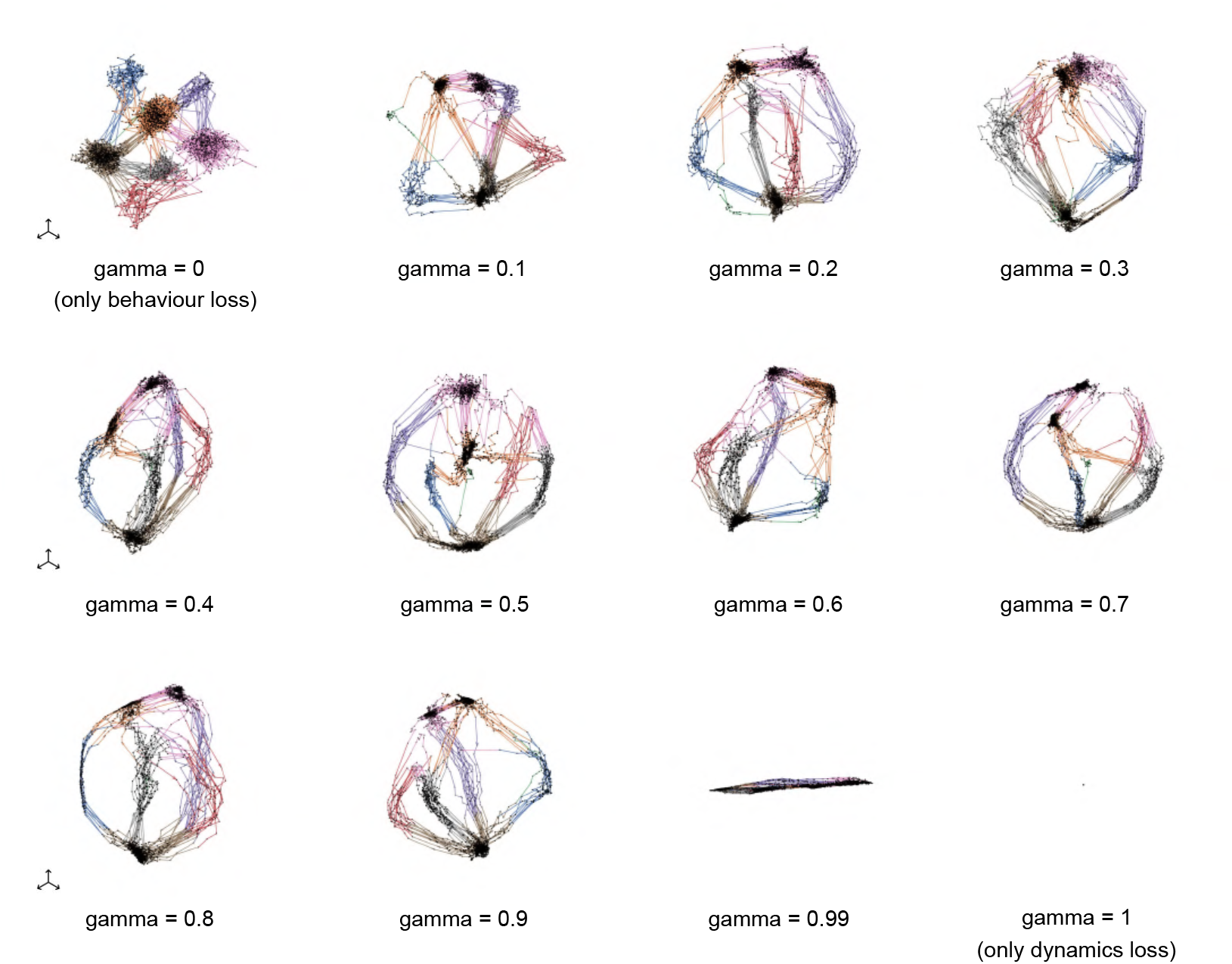
Effect of varying the hyperparameter gamma on the manifold shown on worm-1. The manifolds are plotted in three dimensional latent space.

### S5 Learning curves on original versus information-ablated data

**Figure S4:**
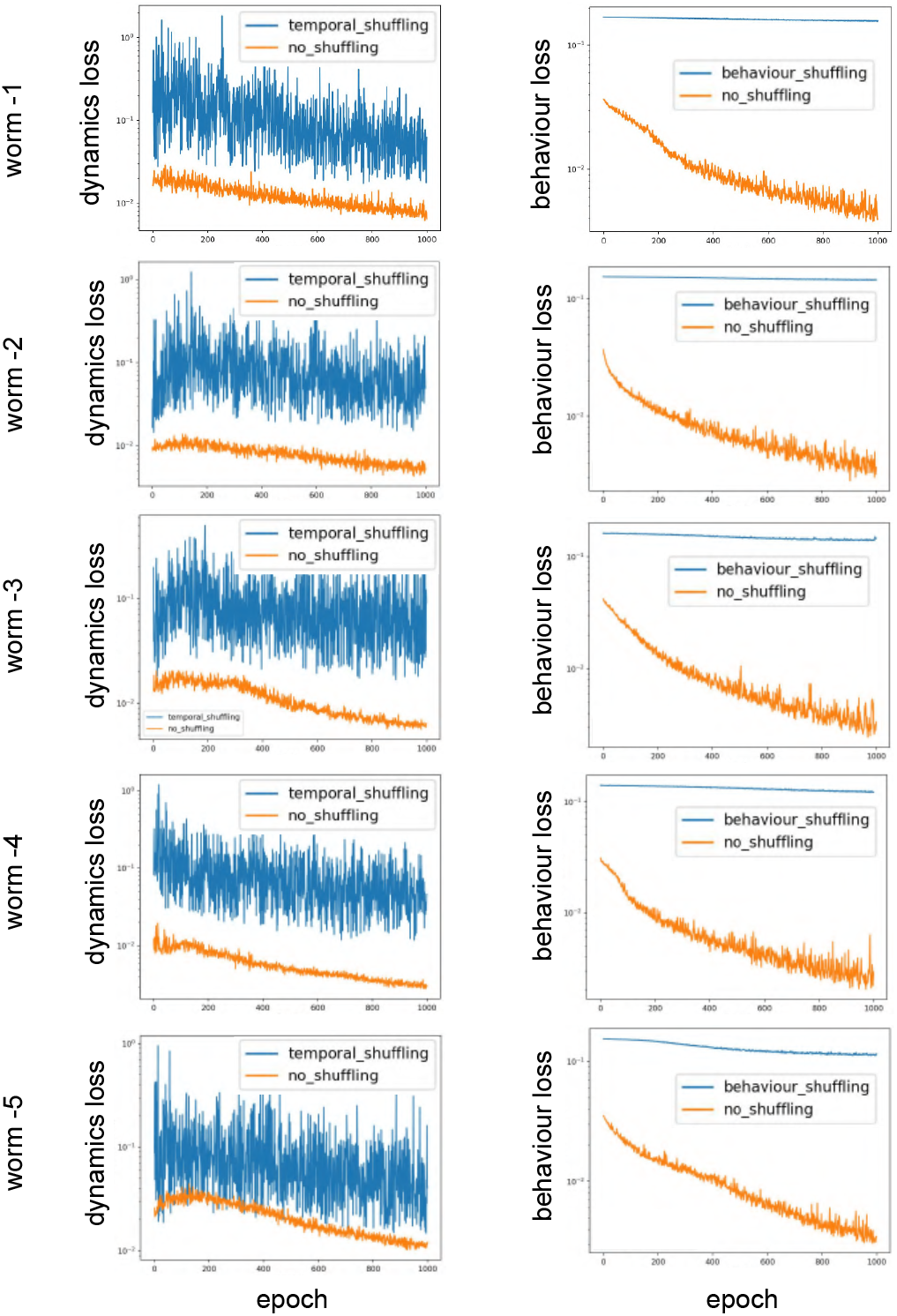
Learning curves on original versus information-ablated data. Learning curves of the dynamics and behaviour losses are shown for all five worms, comparing training on original data with training on shuffled controls (dynamics shuffled or behaviour labels shuffled).

On original data, losses decrease rapidly, whereas on shuffled data they remain nearly flat, indicating that the model extracts meaningful dynamical and behavioural structure rather than fitting noise. These comparisons show that the relevant information is indeed present in the recordings and that the model can capture it. Moreover, this approach provides a diagnostic: if learning curves on original and shuffled data were indistinguishable, either the data lacks the relevant information or the model fails to capture it. Similarly, if the model were able to fit shuffled data, it would signal overfitting. This control establishes that the BunDLe-Net manifold reflects meaningful behavioural and dynamical information inherent in the data, and not overfitting or noise.

### S6 Further evaluation of BunDLe-Net’s embedding

**Figure S5:**
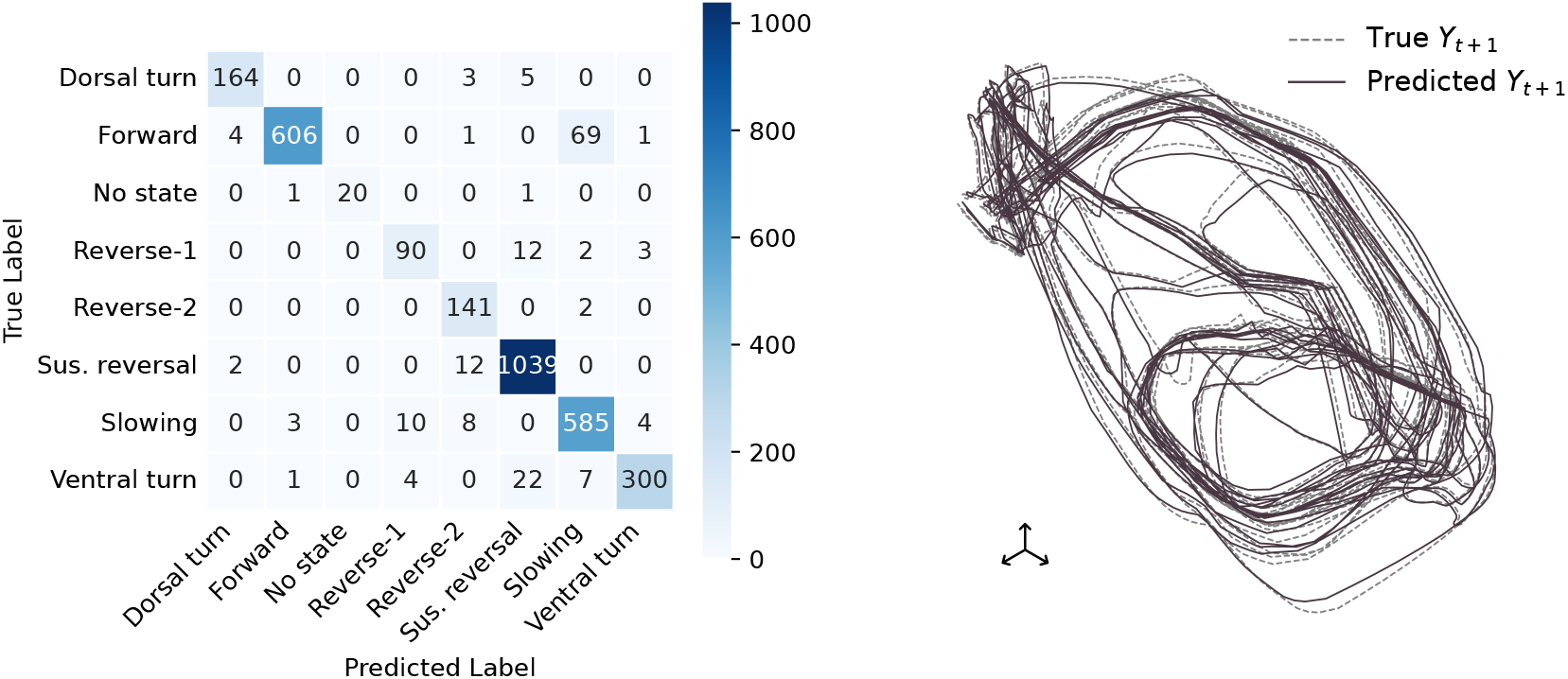
(left) Confusion matrix of behaviour predictions from BunDLe-Net’s embedding for Worm-1 (right) True dynamics and dynamics predicted by BunDLe-Net for Worm-1.

In the following, we provide further information to build an intuition for the behavioural and dynamic prediction performance of BunDLe-Net. In Fig. S5 a), we present the confusion matrix for BunDLe-Net’s behavioural *prediction layer* from the ANN architecture. BunDLe-Net achieves a decoding accuracy of 94.3%, with the few decoding errors dominated by confusion of forward and slowing, two behaviours that are qualitatively similar and only quantitatively differ in the speed of the motion. To evaluate the dynamical performance of the model, we use the *transition model layer T*_*Y*_ to estimate *Y*_*t*+1_ from *Y*_*t*_ and compare it with the true *Y*_*t*+1_, obtained as *τ* (*X*_*t*+1_). Fig. S5 b) shows that the predicted dynamics indeed track the true dynamics rather well. These results indicate that the behaviour predictor and transition model within BunDLe-Net do well to preserve dynamical and behavioural information as intended.

### S7 BunDLe-Net architecture

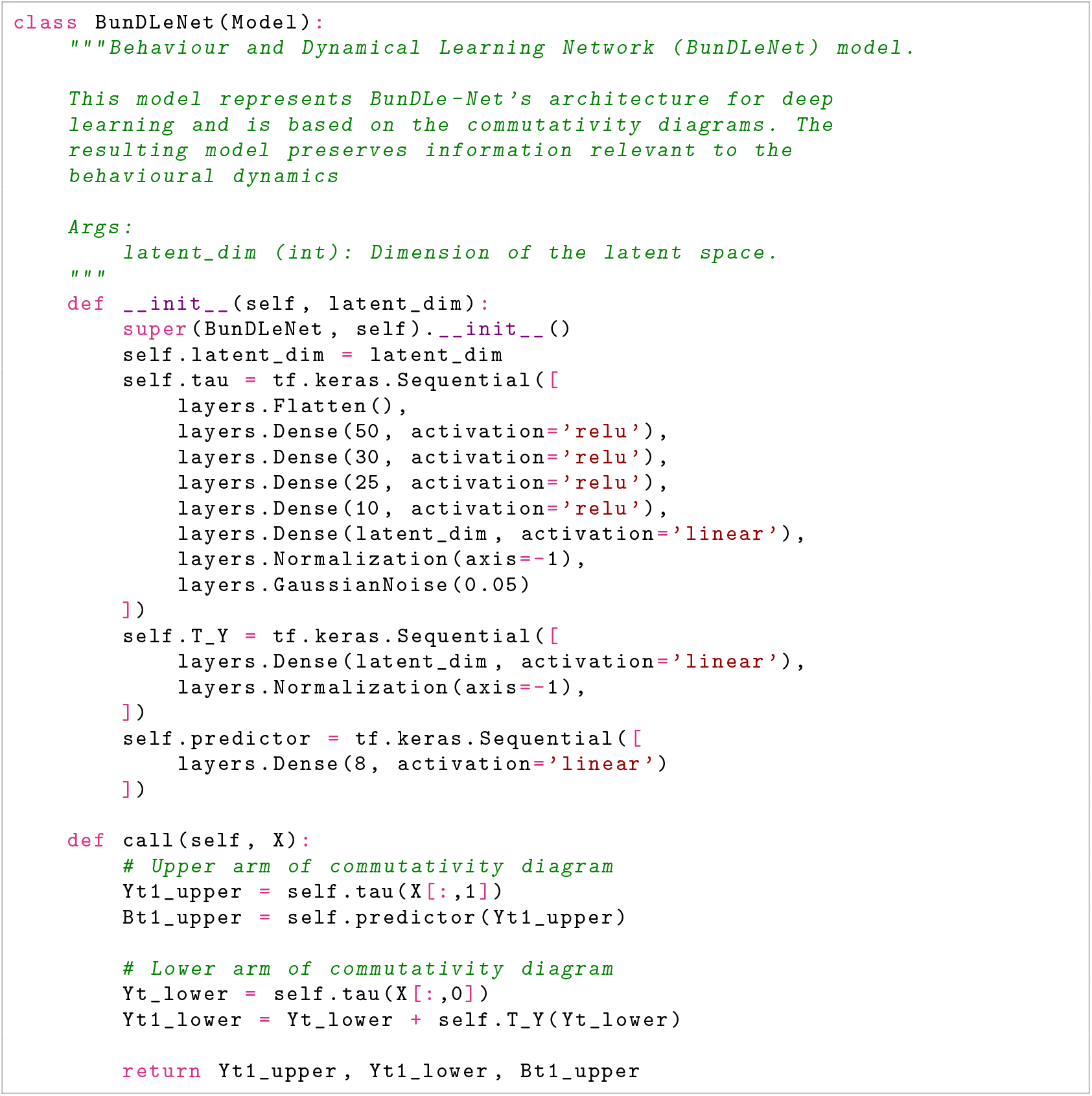

#### S7.1 BunDLe-Net loss function

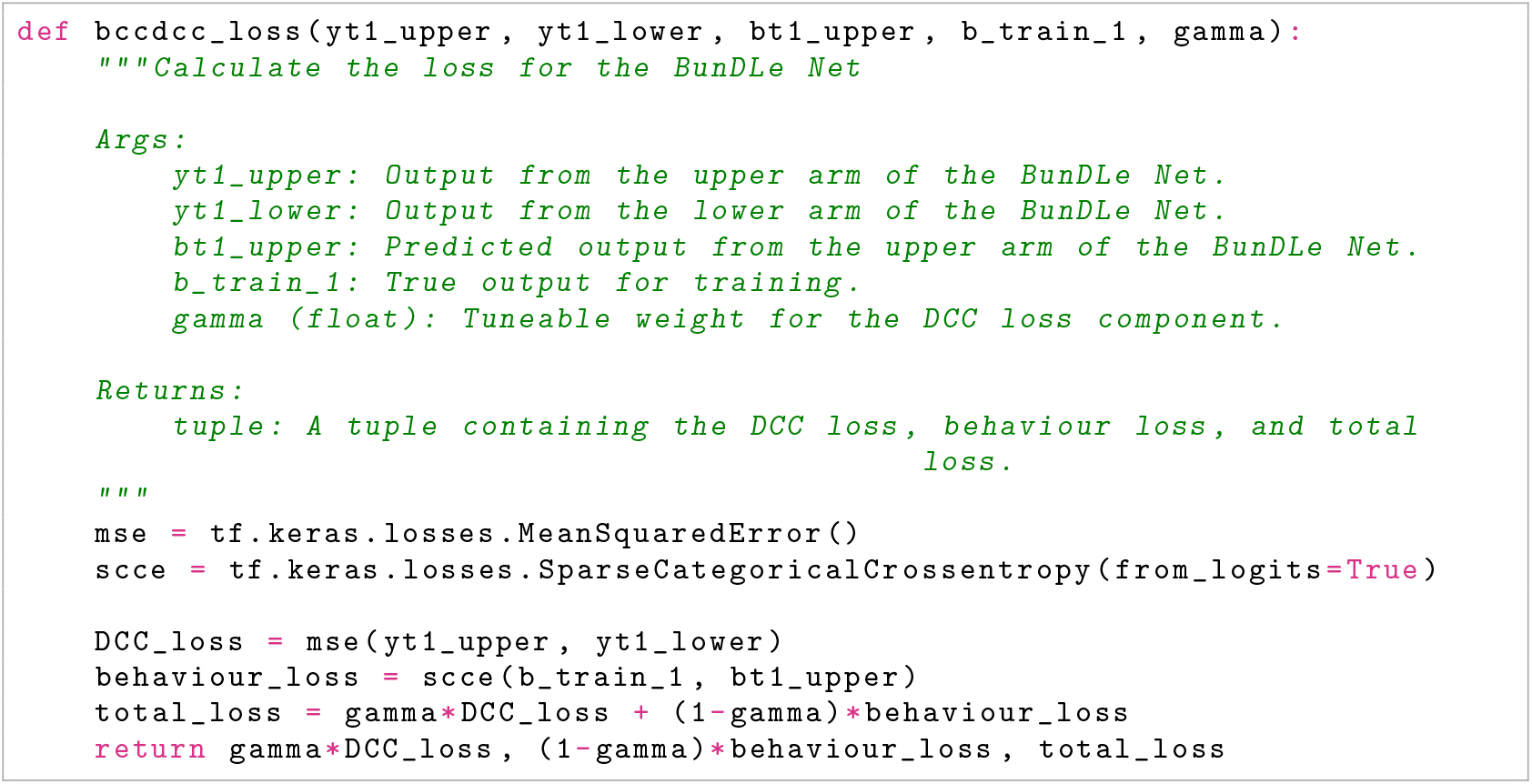

### S8 Embedding of states in distinct behavioural motifs

**Figure S6:**
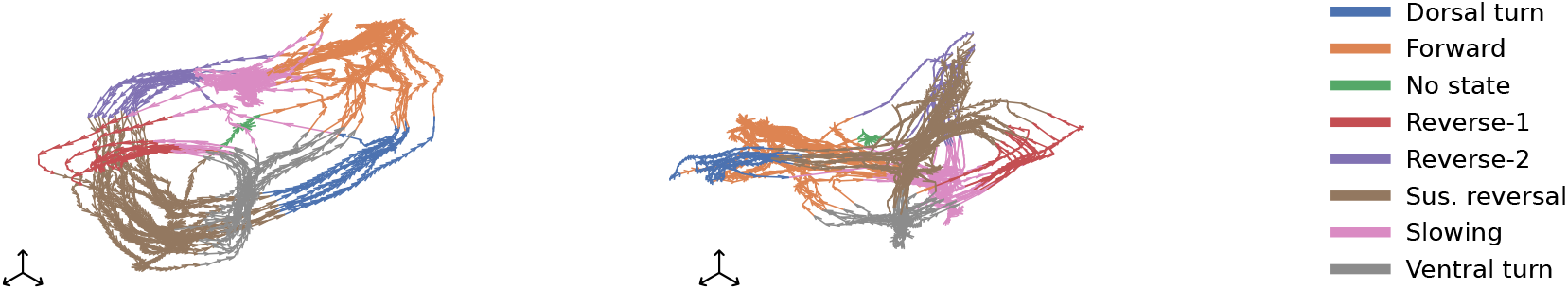
(left) BunDLe-Net trajectory of worm-1 highlighting the embedding of slowing behaviour (in pink) within two distinct bundles or behavioural motifs. (right) In contrast, the sustained reversal state (in brown) is represented by a single intersection point on the right side.

Behaviour can be modelled at different levels of granularity. In the present data set, the worms’ behaviour is described in terms of high-level behavioural patterns such as forward and reversal movements. Alternatively, one could analyse the angular positions and velocities of the various segments of the worms’ bodies, resulting in a more fine-grained representation. Both fine-grained and coarse-grained models hold value in specific contexts. However, it is crucial to maintain consistency within a model’s state space to describe the dynamics accurately. If we utilise a model to understand fine-grained elements but only have access to coarse-grained information, the resulting model will be incomplete or inconsistent in the sense that it lacks the essential information required to predict features of the behavioural dynamics at the desired level of granularity. Here, we demonstrate how BunDLe-Net adeptly handles the coarse-graining of data while still preserving the crucial distinctions between states that are instrumental in explaining the overall dynamics.

We present the discovery of two distinct behavioural states with identical labels based on BunDLe-Net’s neural embedding concerning the given set of behaviours. Consider branches 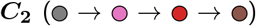 and 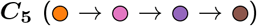 of Fig. 4b. The *slowing* behaviour (in pink) occurs in both these branches, i.e., they are represented distinctly in the latent space and are not fused together even though they have been assigned the same behavioural label. Branch ***C***_**2**_ has a much shorter *slowing* segment than branch ***C***_**5**_. We name the new behavioural states corresponding to ***C***_**2**_ and ***C***_**5**_ as *slowing 1* and *slowing 2*, respectively. These different types of slowing movements are embedded in distinct behavioural motifs since they differ in their neural realisation and their relevance for the model dynamics, i.e., one would predict different future trajectories depending on whether the state is *slowing 1* or *slowing 2*. We note that this is not the case for other behavioural states, e.g., the sustained reversal (in brown), for which all trajectories form one coherent bundle in the embedding. This implies that, in the behavioural state of a sustained reversal, BunDLe-Net found no information at the neural level to predict whether a dorsal or ventral turn is more likely to occur next. In summary, BunDLe-Net can maintain distinct representations or fuse trajectories depending on whether dynamical information about future behaviours is present. Accordingly, if provided with a set of behaviours that are not consistent or complete for the construction of a full dynamical model, BunDLe-Net can discover extra distinctions or *states* that complete this set of behaviours, provided this information is present in the neural activity.

### S9 Learning process

**Figure S7:**
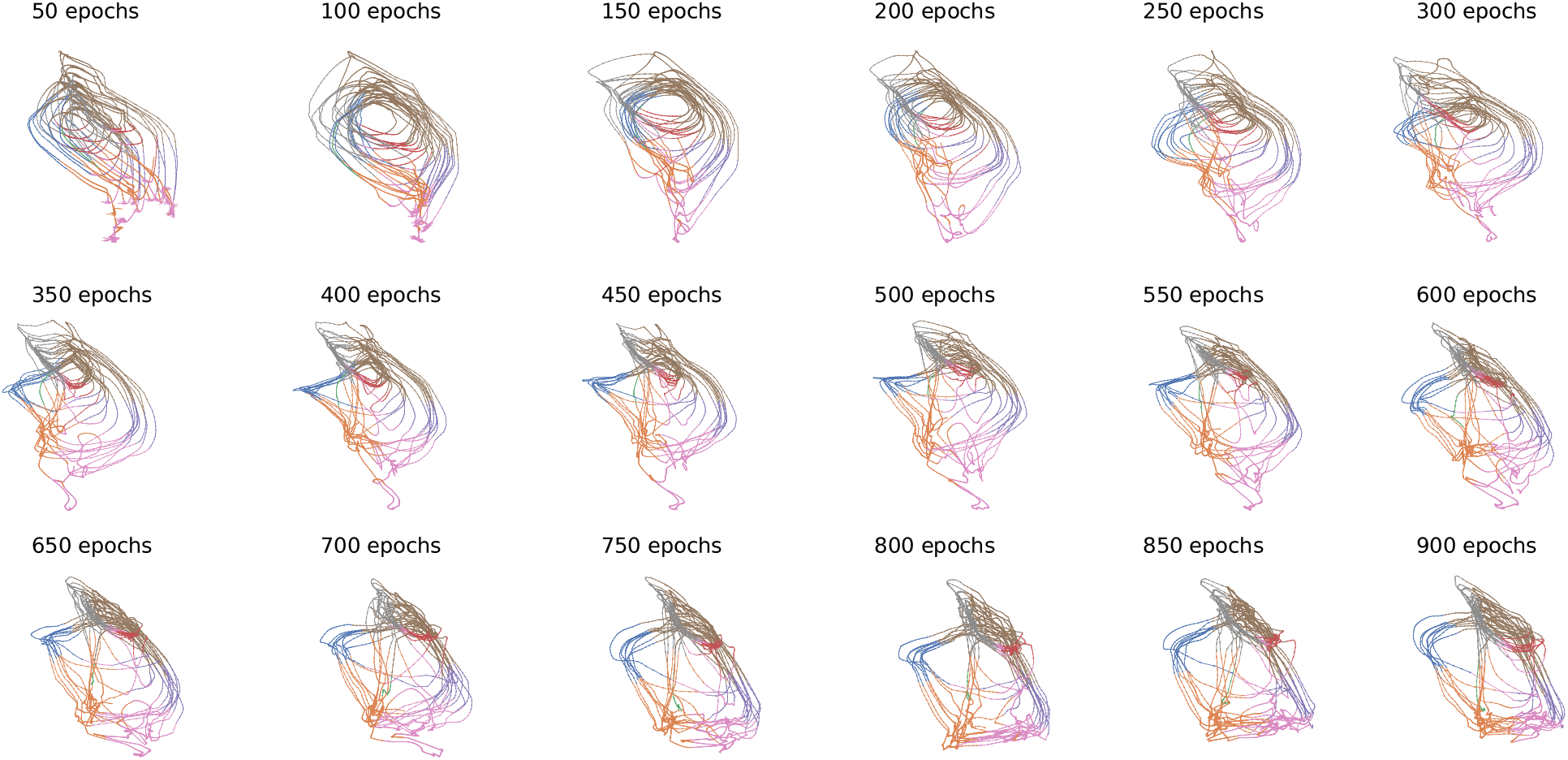
Visualisation of the BunDLe-Net learning process on worm-1 as a function of epochs

### S10 Model ablation studies on rat data

**Figure S8:**
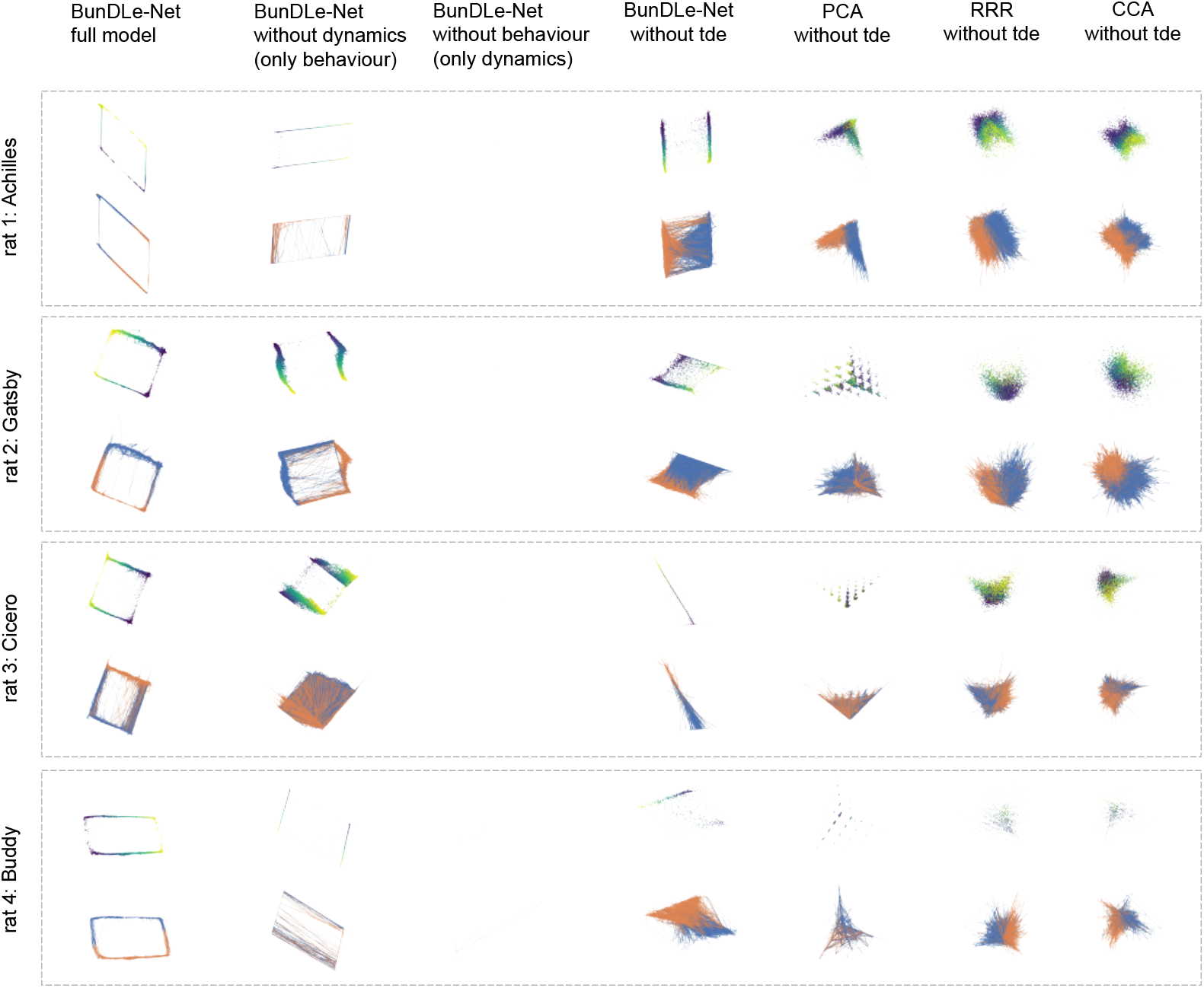
In silico ablation studies on BunDLe-Netand other models on rat data. Data from four rats are shown in separate grey dashed boxes, with columns corresponding to different ablations performed on embedding methods. For each method, the same embedding is shown twice, coloured by position (continuous colour map) and movement direction (binary colour map), respectively, illustrating how behaviour is represented in the latent space. The top row shows only the embedded data points, while the bottom row shows arrows connecting successive time points in order to visualise how the trajectories evolve over time.

In order to see how each feature of BunDLe-Net contributes to the precise embedding structure, we perform experiments by fitting ablated versions of our model on the same rat data. For comparision, we include the non-ablated full model in the first column. The second column shows the embedding by BunDLe-Net fitted without a dynamics loss *L*_*D*_ and only a behaviour decoding loss *L*_*B*_. We see that we obtain a different topology than from the full model: instead of a connected loop, the manifold resembles two linear tracks, one for the left and one for the right running direction, with only very sparse data points between these tracks. From the lower plot for each rat, we further see that the arrow diagram is far more unstructured compared to the full model. This is because it did not consider dynamics, and the transitions are all over the place. Only the full model has two clean paths connecting the separate track,s thus closing the loop.

When we turn off the behaviour loss and use only the dynamics loss, the embedding collapses to a single point, as expected. Since there is no behaviour information to preserve, BunDLe-Net abstracts away all information to achieve the point embedding, which achieves a dynamical loss of zero, since it trivially has all information about its own future at every time point. Next, we consider BunDLe-Net with both losses but without the time delay embedding. That is, instead of providing a window of time steps to the model, we provide only a single time slice. The results in the fourth column show a degradation in the structure of the manifold. This indicates that there is much less information in a single time slice than a history of time slices about both the behaviour variable (position and direction), as well as the dynamics. This aligns with Takens theorem that says that the history of observations allows us to reconstruct the state of a dynamical system even with only partially observed data (Strogatz, 2018; Takens, 1981). Confirming this, in columns five to seven of Fig. S8, we observe a degradation in the topological structure of the embeddings of PCA, CCA, and RRR when the time delay embedding is removed, compared with the corresponding versions that include the time delay embedding in Fig. 8.

### S11 Primate cortex data

**Figure S9:**
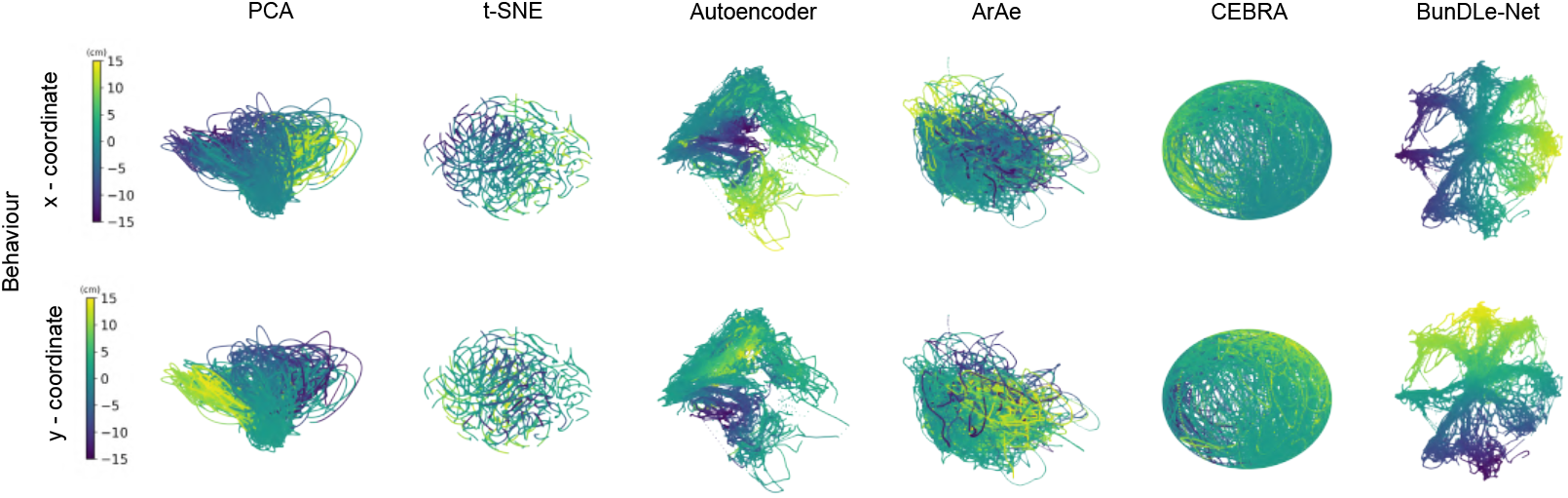
Embeddings in 3-D space of primate neural data.

#### S11.1 Dataset description

The data comprises neural recordings from Area 2 of the primary somatosensory cortex of a primate engaged in a centre-out eight-direction reaching task. This dataset, initially presented by Chowdhury et al. (Chowdhury et al., 2020), is publicly available and has served as a benchmark for neural manifold learning (Schneider et al., 2023; Zhou and Wei, 2020b). We used pre-processed data from (Schneider et al., 2023), where time bins were employed and the estimated spikes of individual units were convolved using a Gaussian kernel to derive estimates of time-varying average spike rates. The behavioural variable comprises the *x* and *y* coordinates representing the position of a cursor controlled by the monkey during the reaching task. Simultaneously, extracellular neural recordings from the primary somatosensory cortex were captured.

#### S11.2 Embeddings

We now demonstrate BunDLe-Net using data collected from the primate somatosensory cortex during a center-out eight-direction reaching task (Chowdhury et al., 2020). Fig. S9 shows the 3-D latent space embeddings of the neural activity by several algorithms. The colour labelling signifies the x and y coordinates of hand position. From the embedding by PCA and the autoencoder, we see that behavioural information is preserved to some extent since different behavioural states occupy distinct regions of 3-D space in the embedding. This would allow us to decode behaviour from these embeddings. Note, however, that in the autoencoder embeddings, dissimilar behaviours are mapped near each other while similar ones are separated. The violet and yellow regions are located near each other, even though they correspond to positions that are the furthest apart. On the other hand, some green regions are fragmented and interspersed with dissimilar behaviours (lower plot). BunDLe-Net’s embedding on the other hand, demonstrates both a high behaviour decodability while also respecting the similarity structure of the behaviour. In fact, it is the only embedding where the task of reaching out in eight directions is evident from the topology^5^. The primate data embedding is different in structure from the worm and rat embeddings in that it does not have a recurrent structure to the dynamics, but instead a clear beginning and end of the reaching action. In BunDLe-Net’s embedding, we see the beginning of the action depicted by the centre of the embedding, where there is no information about which direction the monkey is going to reach out to. Over time, we see this information appearing as the embedding branches off into one of its eight arms, each of which corresponds to a direction in which the monkey reaches out.

## Footnotes

1 To be more mathematically precise the LHS of the following expression would be *P* (*X*_*t*+1_ ∈ *τ*^−1^(*y*)|*X*_*t*_ = *x*^*′*^). However, since this is less readable, we shall use *P* (*Y*_*t*+1_ = *y*|*X*_*t*_ = *x*^*′*^) as shorthand for it. This is possible since *Y*_*t*+1_ = *y* and *X*_*t*+1_ ∈ *τ*^−1^(*y*) are equivalent statements if *τ* links their spaces. Thus, the LHS lives in the probability space of *X* while the RHS lives in the probability space of *Y*.

2 *Note*: This proof is done for the case of discrete state space *X*. The statement is also thought to hold for continuous state spaces, but the proof would be considerably more involved and is hence omitted.

3 Note that matrix representations in the probability space [0, 1]^|*X*|^ and [0, 1]^|*Y*|^ do not necessarily mean linear-functions in the state space *X* and *Y*. Thus, we are not making any restrictions on *T*_*X*_ and *τ*, which are free to be non-linear on the state spaces

4 mapping to a lower-dimensional space

5 No post-hoc averaging was required to reveal the behavioural patterns in the embedding, unlike other methods as in (Schneider et al., 2023)

